# Predicting which genes will respond to perturbations of a TF: TF-independent properties of genes are major determinants of their responsiveness

**DOI:** 10.1101/2020.12.15.422864

**Authors:** Yiming Kang, Michael R. Brent

## Abstract

**Background:** The ability to predict which genes will respond to perturbation of a TF’s activity serves as a benchmark for our systems-level understanding of transcriptional regulatory networks. In previous work, machine learning models have been trained to predict static gene expression *levels* in a given sample by using data from the same or similar conditions, including data on TF binding locations, histone marks, or DNA sequence. We report on a different challenge – training machine learning models that can predict which genes will respond to perturbation of a TF *without using any data from the perturbed cells*.

**Results:** Existing TF location data (ChIP-Seq) from human K562 cells have no detectable utility for predicting which genes will respond to perturbation of the TF, but data obtained by newer methods in yeast cells are useful. TF-independent features of genes, including their pre-perturbation expression level and expression variation, are very useful for predicting responses to TF perturbations. This shows that some genes are poised to respond to TF perturbations and others are resistant, shedding significant light on why it has been so difficult to predict responses from binding locations. Certain histone marks (HMs), including H3K4me1 and H3K4me3, have some predictive power, especially when downstream of the transcription start site. In human, the predictive power of HMs is much less than that of gene expression level and variation. Code is available at https://github.com/yiming-kang/TFPertRespExplainer.

**Conclusions:** Sequence-based or epigenetic properties of genes strongly influence their tendency to respond to direct TF perturbations, partially explaining the oft-noted difficulty of predicting responsiveness from TF binding location data. These molecular features are largely reflected in and summarized by the gene’s expression level and expression variation.

## INTRODUCTION

Understanding the function of a genome requires knowing which transcription factors (TFs) directly regulate each gene. A systems-level understanding should also enable us to predict which genes will change in expression level in response to direct perturbations of TFs. It was hoped that determining where in the genome each TF binds by chromatin-immunoprecipitation (ChIP) would go a long way toward solving these problems, but several studies have shown that the set of genes whose promoters are bound by a TF and the set of genes that respond to perturbations of that TF do not overlap much (Gitter et al. 2009; Lenstra and Holstege 2012; Cusanovich et al. 2014; Kang et al. 2020). Genes that are responsive but not bound may be indirect targets of the TF. The genes that are not responsive despite the fact that their regulatory DNA is bound by the perturbed TF constitute a greater mystery. Currently, we cannot predict which bound genes will respond to a perturbation and which will not. In this paper, we take on the challenge of predicting whether a gene will respond to perturbation of a TF by using data on where the TF binds along with a variety of TF-independent features of each gene, including histone marks (HMs), chromatin accessibility, dinucleotide frequencies, and the gene’s pre-perturbation expression level and expression variation.

A number of studies have shown success in predicting the expression *levels* of genes by using TF binding signals (Middendorf et al. 2004; Ouyang et al. 2009; Schmidt et al. 2017) or HMs in each gene’s regulatory region (Karlić et al. 2010; Cheng et al. 2011; Dong et al. 2012; McLeay et al. 2012; Singh et al. 2016; Read et al. 2019). All these models predict expression level in a given sample by using data from the same cell type and similar growth conditions. As result, the epigenetic features used for prediction could be causes, consequences, or merely correlates of gene expression level (Henikoff and Shilatifard 2011). Recently, deep neural networks have been used to predict the expression level of a gene from the DNA sequence flanking it (Kelley et al. 2018; Zhou et al. 2018; Washburn et al. 2019; Agarwal and Shendure 2020). Models have also been trained to predict the variability of gene expression within or across cell types (Ouyang et al. 2009; Zhou et al. 2014; González et al. 2015; Crow et al. 2019; Sigalova et al. 2020). In addition to the above genomic features, combining the binding signals of RNA-binding proteins and microRNA at gene bodies with TFBS at promoters was also found to be predictive (Tasaki et al. 2020).

We have taken on a different challenge – training machine learning models that can predict which genes will respond to perturbation of a TF without using any data from perturbed cells. Because the predictive features are measured in unperturbed cells, they cannot be consequences of the perturbation or the response. The overall accuracy of the models serves as a benchmark for our understanding of global regulatory networks. Perhaps more important, analysis of the trained models can provide insight into the factors that determine which genes respond to a TF perturbation. Many methods have been developed to explain how specific features and feature values influence a complex model’s predictions (Molnar 2019; Breiman 2001; Zeiler and Fergus 2012; Zhou and Troyanskaya 2015; Fisher et al. 2019). In this paper we rely on SHAP values (Lundberg and Lee 2017). A SHAP value represents the influence of one feature on one prediction. The SHAP value for a given prediction is based on a locally weighted linear regression model that approximates a non-linear model in the region of feature space near that prediction. A positive SHAP value for a feature of a prediction indicates that that feature pushes the model to predict a higher response. Conversely, a negative SHAP value indicates that the feature value pushes the model to predict a lower response. The magnitude of the SHAP value of a feature of a prediction indicates how much that feature influences that prediction.

Summary calculations make it possible to draw conclusions about features that apply to all predictions or to a specific subset of prediction. Separately summing the positive and negative SHAP values for a feature over a set of predictions reveals the relative strength of the positive and negative influences of the feature. This can be especially useful when looking at just the positive predictions, just the negative predictions, predictions which are predicted accurately or inaccurately. Summing positive and negative SHAP values of a feature together provides a sense of the feature’s overall direction of influence, which we refer to as its *net influence*. Summing the absolute values of the SHAP values for a set of predictions shows how important the feature is in determining the model’s predictions, regardless of direction. We refer to the sum of absolute SHAP values over a set of predictions as *global feature importance*.

SHAP analysis, complemented by analyses of model accuracy, provides several surprising biological and methodological insights.

1. Existing genome-scale data on TF binding locations, including ENCODE data on human K562 cells, are not useful for predicting which genes will respond to perturbation of a TF. However, yeast data obtained by newer methods (transposon calling cards or ChIP-exo) are.
2. A few HMs have value for predicting perturbation responses, primarily when they occur in the gene body downstream of the transcription start site (TSS).
3. For both yeast and human, pre-perturbation gene expression level and gene expression variation (GEX features) were surprisingly useful for predicting whether a gene would respond to perturbation of any TF or other regulatory protein; for human cells, they were far and away the most useful features. When these features are available, HMs provide no additional information that is useful for predicting perturbation responses in human K562 cells.

In summary, properties of the gene itself have a major influence on its tendency to respond to regulatory perturbations. The extent to which this tendency is determined by the gene’s epigenetic state or its inherent properties remains to be seen.

## RESULTS

### Modeling frameworks, features, and datasets

We took a two-step approach to understanding the determinants of transcriptional responses to TF perturbations: (1) train machine learning models to predict whether each gene will respond to a perturbation of a particular TF and (2) analyze the trained models to identify which genomic features they used to make their predictions. We provided the models with three types of genomic features (Fig. 1A). First, data on the binding locations of the perturbed TF (location features). Second, data on the median and variance of each gene’s expression levels in unperturbed samples (GEX features).

**Figure 1.**
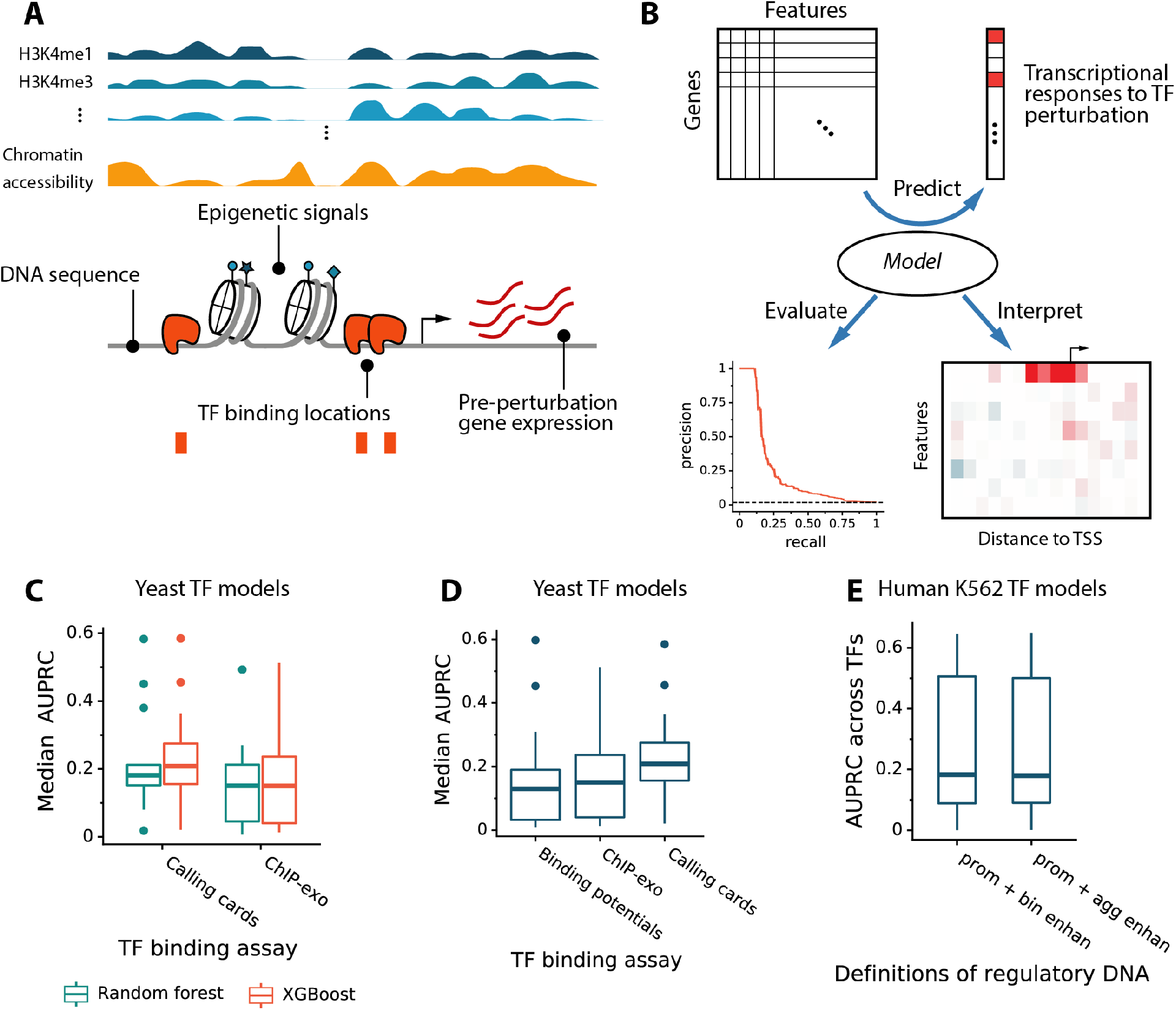
Model and performance. (A) Features for predicting transcriptional responses to TF perturbation. (B) Framework for predicting responses, evaluating model performance, and estimating local feature influences. (C) Model accuracy on yeast TFs using all binding location data. (D) Model accuracy on eight yeast TFs with binding location data from both ChIP-exo and calling cards assays. (E) Model accuracy on human K562 cells using two methods of aggregating data from enhancers associated with each gene.

Third, data on each gene’s epigenomic context, including DNA accessibility, selected histone modifications, and dinucleotide frequencies (epigenetic features). We focused on eight histone marks that were previously shown to be most useful for predicting gene expression level (Karlić et al. 2010; Zhou et al. 2014; González et al. 2015; Singh et al. 2016; Roadmap Epigenomics Consortium et al. 2015) (Table S1). Neither GEX features nor epigenetic features are tied to any specific TF – if they predict a gene’s responsiveness to perturbation of one TF, they should also predict its responsiveness to perturbation of other TFs. To generate a feature matrix, we defined cis-regulatory regions for each gene and mapped genomic data to them. For yeast genes, we assumed a regulatory region ranging from 1000 bp upstream of the transcription start site (TSS) to 500 bp downstream. Although most studies assume the yeast promoter is smaller than this, we expected that the models would learn which parts of this region are most predictive. For human genes, we included both proximal promoters (4 kb centered on the 5’-end TSS) and distal enhancers (taken from (Fishilevich et al. 2017); see Methods). 47% of alternative TSS’s fell within 4 kb region around the 5’ TSS. The 5’ TSS and others within 2 kb of it account for the vast majority of transcription (Supplemental Fig. S1). Alternative promoters outside of this region were treated as enhancers (Andersson and Sandelin 2020). To test whether certain locations within a regulatory region are more important than others, we divided the promoter regions into 100 bp subregions, each with its own features. Within each 100 bp subregion, signals from assays for TF binding location, DNA accessibility, or histone marks were aggregated and discretized. For yeast, we used TF binding location data generated by two *in vivo* assays: transposon calling cards (Wang et al. 2011; Shively et al. 2019; Kang et al. 2020) and ChIP-exo (Bergenholm et al. 2018; Rossi et al. 2018). We showed previously that these datasets predict perturbation responses much better than older ChIP-chip data (Kang et al. 2020). We used data on yeast histone marks from ref. (Weiner et al. 2015) and chromatin accessibility from ref. (Schep et al. 2015), both assayed in steady-state growth conditions. For human models, we used data from the K562 cell line because the it has the most TFs that were ChIPped and perturbed in the ENCODE Project (Dunham et al. 2012; Davis et al. 2018; Abascal et al. 2020). Histone marks, DNA accessibility, and perturbation-response data were also from ENCODE (see Methods). For both yeast and human, preperturbation expression variance was adjusted to make it independent of expression level (Methods; Supplemental Fig. S2).

We trained the models to predict whether a gene will respond to a TF perturbation (Fig. 1B). For yeast, responsiveness was determined by using data from ref. (Hackett et al. 2020), who measured transcriptional responses shortly after chemically inducing overexpression of each TF. We focused on the responses at 15 minutes after the induction (see Methods). For human, responsiveness was determined by using ENCODE RNA-Seq data measured after TF knockdown or knockout. Our datasets included 25 yeast TFs and 56 human TFs with both binding and perturbation-response data. The average number of genes that responded to each perturbation was ~6.7% in yeast (median: 2.7%, sd: 9.4%) and ~6.4% in human (median: 2.5%, sd: 8.3%).

We trained and tested two ensemble classifiers for each perturbed TF—random forests and a gradient boosting implementation called XGBoost (Chen and Guestrin 2016) – by using ten-fold cross-validation on genes. Below, we analyze how the features influence the prediction for each gene using the model that was not trained on that gene. We used precision recall curves for accuracy evaluation and the area under the curve (AUPRC) as a summary statistic. This approach is appropriate because only a small fraction of genes is responsive to each perturbation, creating large class imbalances.

First, we tested the two classifiers using yeast TF binding-location data from either transposon calling cards or ChIP-exo, keeping all other features constant. The best combination of classifier and binding data was XGBoost on calling cards data (Fig. 1C). However, calling cards and ChIP-exo data assayed different sets of TFs. To make a direct comparison, we trained and tested XGBoost models for the eight TFs assayed by both methods. The calling cards data again yielded greater prediction accuracy (Fig. 1D). We also tried replacing binding location data with TF binding potentials obtained by scanning a binding specificity model for the perturbed TF (Spivak and Stormo 2012; Grant et al. 2011) over promoter sequences. Binding potential was least useful, even when data on chromatin accessibility was also included in the model (Fig. 1D). Going forward, we use the XGBoost models. For yeast, we focused on TFs that had either calling cards or ChIP-exo data; for those that had both, we used the data that yielded the best prediction accuracy.

Using XGBoost on the K562 ENCODE data, we investigated two ways of incorporating binding and epigenetic features from enhancers. The two methods divide the region around the promoter into subregions in different ways and sum the signals from enhancers within each subregion to form a single feature value. The first method (*bin enhan*) sums signals over enhancers within subregions whose widths increase exponentially with their distance from the TSS (Supplemental Fig. S3). The second approach (*agg enhan*) sums signals from all enhancers upstream of the TSS to create one feature and all enhancers downstream of the TSS to create another. Models trained using the two strategies of enhancer-feature mapping show no significant difference in accuracy (P = 0.63, paired t-test; Fig. 1E), so we used less numerous aggregated enhancer features in the remainder of the study.

The prediction accuracy varied quite a bit from one TF to another (Supplemental Fig. S4). In general, accuracy was lower for TFs that had fewer responsive targets than for those that had many (Supplemental Fig. S5A,B). This is likely the result of extreme class imbalance, which is known to hinder classification algorithms (Japkowicz and Stephen 2002) and the lack of enough positive examples to learn from. In the ENCODE data, another major factor was the effectiveness of the TF perturbation. The larger the absolute log fold change of the TF in the perturbed sample relative to the unperturbed, the better the TF model performed (Supplemental Fig. S5D). There was no such trend in the yeast data because the induced TFs were highly over-expressed (typically at least 16-fold, Supplemental Fig. S5C).

### SHAP analysis shows that the TF binding signal is useful for prediction in yeast

The next step in our analysis was to determine what XGBoost learned about genomic features and how it used them. The model interpretation approach we used is based on SHAP values (Lundberg and Lee 2017; Lundberg et al. 2018). SHAP values explain why the prediction for one particular test example – one TF-gene pair – differs from the average prediction for all genes in response to perturbation of that TF. Of course, explaining what an algorithm learned is only interesting if it learned something significant, as indicated by its prediction accuracy. Thus, we analyzed only models with a median cross-validation AUPRC greater than 0.1, yielding models for 17 yeast TFs (Supplemental Table S2) and 30 human TFs (Supplemental Table S3).

Taking the XGBoost model for yeast TF Lys14 as an example, Figure 2A illustrated the SHAP values calculated for each feature of *LYS9*, a responsive gene, and *ECM23*, a non-responsive gene. The primary factors that caused the model to predict that *LYS9* would be responsive are (1) the Lys14 binding signal in the 500 bp upstream of the TSS and (2) *LYS9*’s pre-perturbation expression level (Fig. 2A, left, red). For *ECM23*, these positive influences were absent (Fig. 2A, right). Furthermore, *ECM23*’s pre-perturbation expression level and, to a lesser extent, its pre-perturbation expression variation, pushed the model to predict that it would not respond to Lys14 perturbation (blue). To aggregate these influences across promoter regions, we separately summed all positive SHAP values for each feature, which are plotted in red to the right of the heatmaps, and all negative SHAP values for each feature, which are plotted in blue. If a feature has a positive influence in some regions of the promoter but negative in others, it is shown with both a red bar (to the right of the centerline) and a blue bar (to the left of the centerline). For *LYS14* and *ECM23*, most features have only positive or only negative SHAP values, so only one bar is visible.

**Figure 2.**
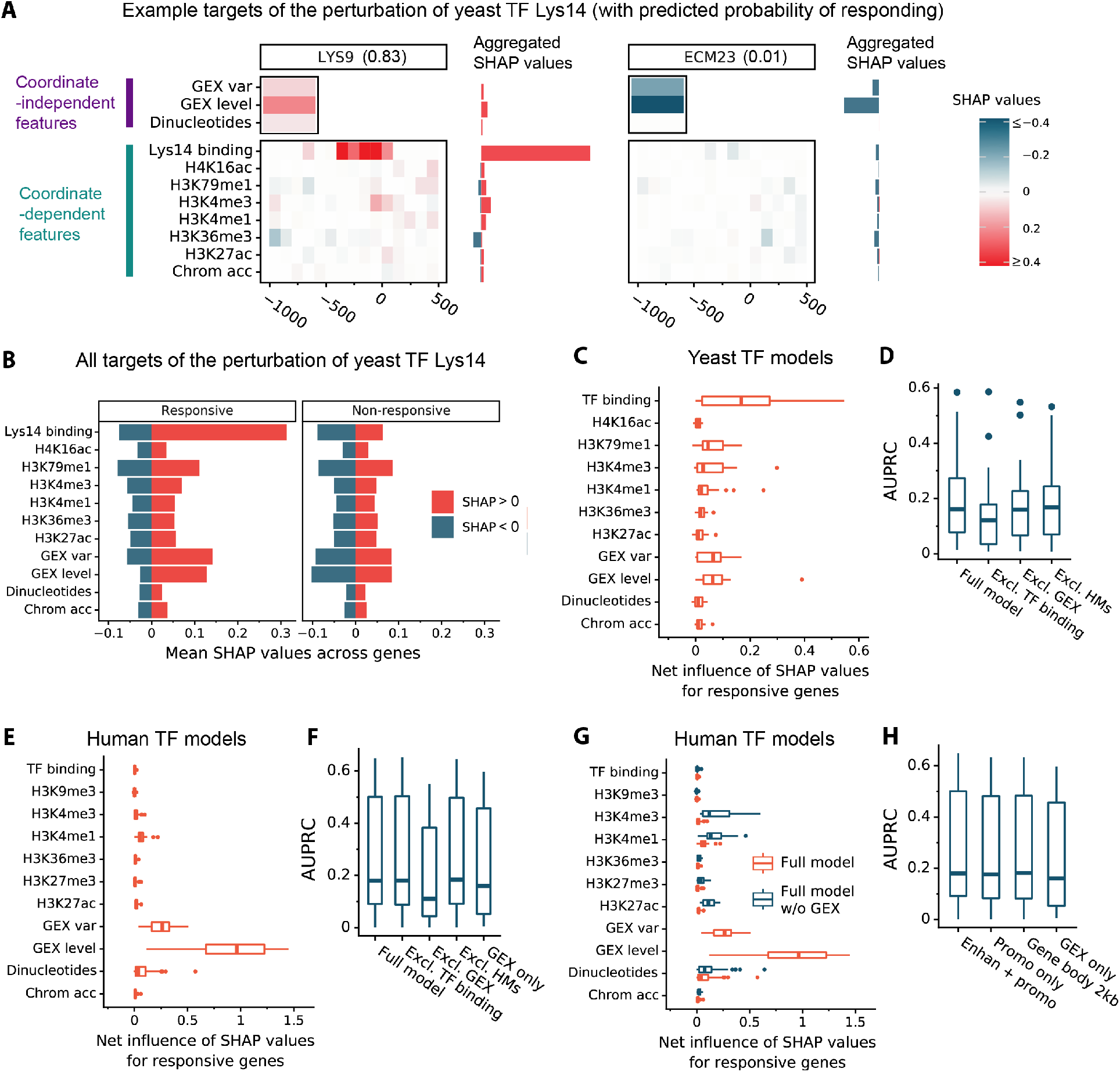
Quantification of feature influences. (A) An example of decomposing the predicted score using SHAP values. Lys9 is a responsive target of yeast TF Lys14 with predicted response probability 0.83 and Ecm23 is an unresponsive gene with predicted response probability 0.01. The top panel shows the features that are independent of genomic coordinates; the bottom panel shows the features that depend on genomic coordinates. The right horizontal bars show the respective sums of SHAP values that are positive (red) and negative (blue), regardless of their genomic coordinates. (B) Left: For yeast TF Lys14, the positive (red) or negative (blue) SHAP values for each feature, summed over genomic positions relative to each gene and averaged over genes that respond to Lys14 perturbation. Right: The same analysis for genes that do not respond to Lys14 perturbation. (C) Distribution across TFs of the “net influence” of each feature on predictions, averaged over responsive targets. Net influence is the sum of all SHAP values for a feature, regardless of sign or genomic position. (D) Comparison of yeast model accuracy using four types of input features: the model described previously (*Full model*), the model trained without TF binding features, the model without gene expression features, and the model without histone marks (HMs). (E) Same as (C) except for human K562 TF perturbations. (F) Human model accuracy as in (D), with the addition of a model trained only on gene expression features (*GEX only*). (G) Comparison of net influences of features on predictions for responsive human genes. Models were trained with gene expression features (*Full model*) or without. (H) Comparison of model accuracy using four types of input features: the model described previously (*Full model*), the model excluding enhancer features (*Promo only*), the model excluding enhancer features and features mapped upstream of the TSS (*Gene body 2Kb*), and the model using only pre-perturbation gene expression features (*GEX only*).

To get a sense of how feature values affected the model’s predictions for all genes, we first divided genes into responsive and non-responsive. Within each group, for each feature, we separately summed its positive SHAP values from all promoter regions of all genes and its negative SHAP values (Fig. 2B). For Lys14-responsive genes (Fig 2B, left), Lys14 binding data in the gene’s promoter tends to have a much bigger effect on predictions when it pushes the predicted probability of response up (red bar) than when it pushes the predicted probability of response down (blue bar). Comparing the red and blue bars for other features reveals net positive effects from pre-perturbation gene expression level and variation. Histone marks H3K79me1 and H3K4me3 have smaller positive influences and they have negative influences that are almost as large as the positive ones, on average. Thus, depending on the promoter position, the value of the histone mark feature, and the gene, these features can either increase the predicted probability of a gene’s responding or decrease it. For genes that do not respond to the Lys14 perturbation, the net influences of all features are close to zero (Fig. 2B, right), indicating that they do not push predictions for non-responsive genes very far from the average prediction for all genes. This average is low – 6.8% probability of being responsive – since the vast majority of genes do not respond to perturbation of Lys14.

To generalize from Lys14 to all TFs, we calculated the net influences of features on predictions for genes that respond to perturbation of each TF and plotted the distributions (Fig. 2C). This showed that the findings for Lys14 generalize well to the other TFs. The biggest net influence was the binding signal from the perturbed TF, followed by gene expression level, gene expression variation, and histone marks H3K79me1 and H3K4me3. Supplemental Figure S6 shows the positive and negative influences of each feature on both responsive and non-responsive genes. Complementary analysis of the effects of dropping feature classes from the model confirmed that TF binding features contribute most to the accuracy of the full model, followed by gene expression features (Fig. 2D). Dropping histone marks had a marginally significant but very small effect (the mean AUC dropped 0.01, *P*<0.04). This was due to an effect on a minority of TFs, since the median AUC actually *increased* by 0.006.

### In human cells, ChIP-seq peaks and epigenetic marks have relatively little value for response prediction

Next, we summarized SHAP values for each human TF model, focusing first on genes that respond to perturbation of the TF. Strikingly, ChIP-seq peaks for a TF, which reflect its binding location, had essentially no net influence on predictions for genes that are in fact responsive (Fig. 2E). This is consistent with earlier studies (Gitter et al. 2009; Lenstra and Holstege 2012; Cusanovich et al. 2014; Kang et al. 2020). Gene expression level in unperturbed control samples was the most influential factor, followed by expression variation in the control samples. H3K4me1 and dinucleotide frequencies in the cis-regulatory DNA had very small influences on the predictions for some TF models, but the effects of the other histone marks and of chromatin accessibility were negligible. Analysis of non-responsive genes yielded similar conclusions (Supplemental Fig. S6B). Analysis of predictive accuracy supported these conclusions: Dropping the TF binding locations or the HM features had negligible impact on prediction accuracy, whereas dropping the gene expression features greatly reduced accuracy (Fig. 2F). In fact, a model using only the gene expression features was almost as accurate as the full model.

We hypothesized that HMs may lack influence in the model because the gene expression features summarize any useful information provided by HMs, as well as other aspects of a gene’s epigenetic state. To test this, we trained a model without the gene expression features and analyzed the influence of the remaining features on predictions for genes that are in fact responsive (Fig. 2G). Removing the gene expression features from the model did increase the influence of H3K4me3 and H3K4me1, supporting our hypothesis. However, the model without gene expression features has low accuracy (median AUPRC 0.11), so the predictive value of HMs is small.

These findings drove us to investigate the utility of features mapped to various regions of the cis-regulatory DNA associated with each gene. When we dropped the TF binding signal, histone modifications, dinucleotide frequencies, and chromatin accessibility from the enhancer regions associated with each gene, the effect on prediction accuracy was negligible (mean AUPRC decreased by 0.009, Fig. 2H). We then tried dropping all features from both the enhancers and the promoter regions upstream of the TSS, leaving only the first 2 kb of the gene body. Again, the effect on accuracy was negligible (median *AUPRC increased* by 0.002 relative to the full model while the mean decreased by 0.007). Dropping these genomic features entirely, leaving only GEX features, also had little effect (mean AUPRC decreased by 0.035, or 14% of the full model’s AUPRC). While the TF binding signal and epigenetic features significantly enhanced prediction accuracy in yeast, they had little predictive value in the human data. What predictive value they did have was entirely due to features mapped to the 2 kb downstream of the TSS.

### In yeast cells, TF binding locations and strengths discriminate between bound genes that are responsive and those that are not

The most common use of *in vivo* binding location data is to classify genes into those whose regulatory DNA is or is not bound by the TF. However, this typically yields a large set of genes that are bound by the TF at a statistically significant level but are not responsive to perturbation of that TF (Gitter et al. 2009; Lenstra and Holstege 2012; Cusanovich et al. 2014; Kang et al. 2020). Thus, we investigated whether the model could use the strength and location of the binding signal to better predict which bound genes would be responsive. In Figure 3A, each row shows SHAP values of the TF binding signal in each promoter bin, averaged across the genes that were significantly bound by the perturbed TF. For all but three TFs, the binding signal in the 600 bp upstream of a gene’s TSS influenced the model toward predicting (correctly) that the gene would respond to the perturbation. The Calling Cards and ChIP-exo technologies showed a general concordance on the relative utilities of various positions, but the influence of ChIP-exo was even more localized to 300 bp upstream (Supplemental Fig. S7). Using Leu3 calling cards data as a typical example, stronger binding signals were more influential than weaker ones and signals of the same strength were more influential in the region 100-200 bp upstream of the TSS than in the region 400-500 bp upstream (Fig. 3B). For Leu3, SHAP values in five promoter bins were significantly higher among the bound and responsive group than in the bound but unresponsive group (Fig. 3C). For most TFs, a similar pattern was found in one or more promoter bins (Fig. 3D, Supplemental Fig. S8). Thus, the strength and location of the binding signal are meaningful predictors of whether significantly bound genes will respond to the perturbation, consistent with our earlier findings (Kang et al. 2020).

**Figure 3.**
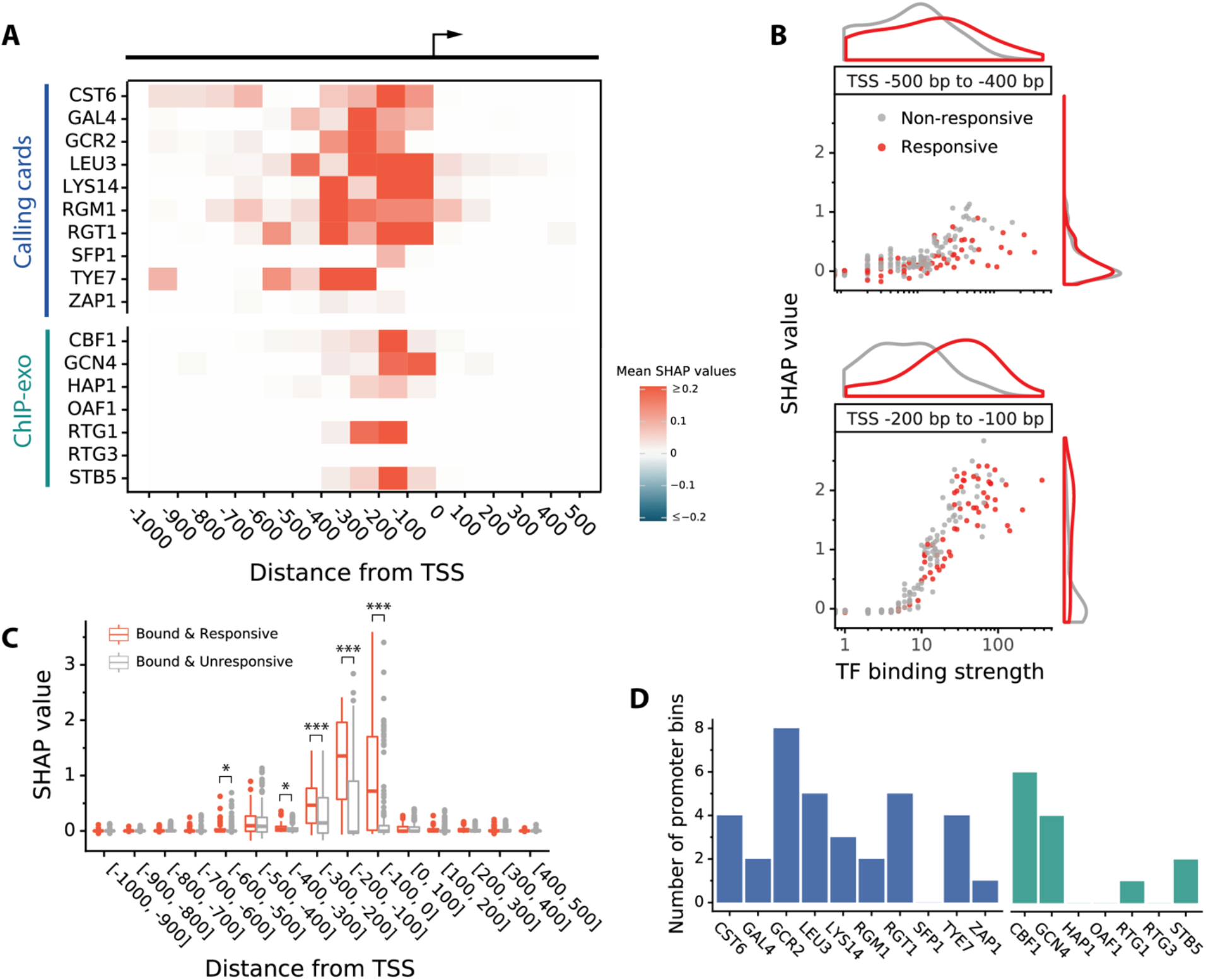
TF binding features in yeast models. (A) Heatmap of the influence of yeast TF binding signals along regulatory DNA. Each pixel is the mean signed SHAP value over all target genes that were bound by the perturbed TFs. (B) Comparison of two upstream bins ([−500, −400] and [−200, −100]) of yeast TF Leu3. Among the genes that are bound by Leu3, the responsive genes are more clearly distinguished from the unresponsive one in the [−200, −100] bin. This shows that even within 500 bp of the TSS, Leu3 binding near the TSS is more likely to be functional than Leu3 binding further away. (C) Comparison of feature influences on responsive and unresponsive targets that were bound by Leu3. *P* < 0.05 (*), p < 0.01 (**), *P* < 0.001(***), Wilcoxon rank-sum test. The significant differences all show that responsive genes are bound more strongly than unresponsive genes. Furthermore, all significant effects of binding strength are within 600 bp upstream of the TSS. (D) Number of 100 bp promoter bins in which the bound and responsive genes have significantly higher SHAP values for TF binding (*P* < 0.05) than the bound but non-responsive genes. Blue bars: calling cards; Green bars: ChIP-exo.

### Highly expressed genes and genes with high expression variation are more likely to be responsive

Given the predictive power of gene expression level and variation, we investigated how the model used these features. Starting with yeast TF Lys14, we noted a monotonic relationship in which the more highly a gene was expressed before the perturbation, the more the model expected it to respond (Fig. 4A). We also noted that, the more a gene’s expression varied from one pre-perturbation sample to another, the more the model expected it to respond (Fig. 4B). This was not due to the relationship between expression level and expression variation, which we removed by fitting a model that predicts expression variation from expression level and using the residuals from that model as our variation feature (Supplemental Fig. S2). For most yeast TFs, both expression level and expression variation are positively correlated with SHAP value – higher expression level and expression variation push the model to predict a higher probability of response (Fig. 4C). The same pattern holds for human TFs (Fig. 4D). The two yeast TFs (Rgt1 and Zap1) and one human TF (ZC3H8) that have negative correlations were those for with the lowest model accuracies, which suggests that either (1) the data on these TFs are not particularly accurate or (2) the model’s feature utilization may not reflect the true patterns in the data.

**Figure 4.**
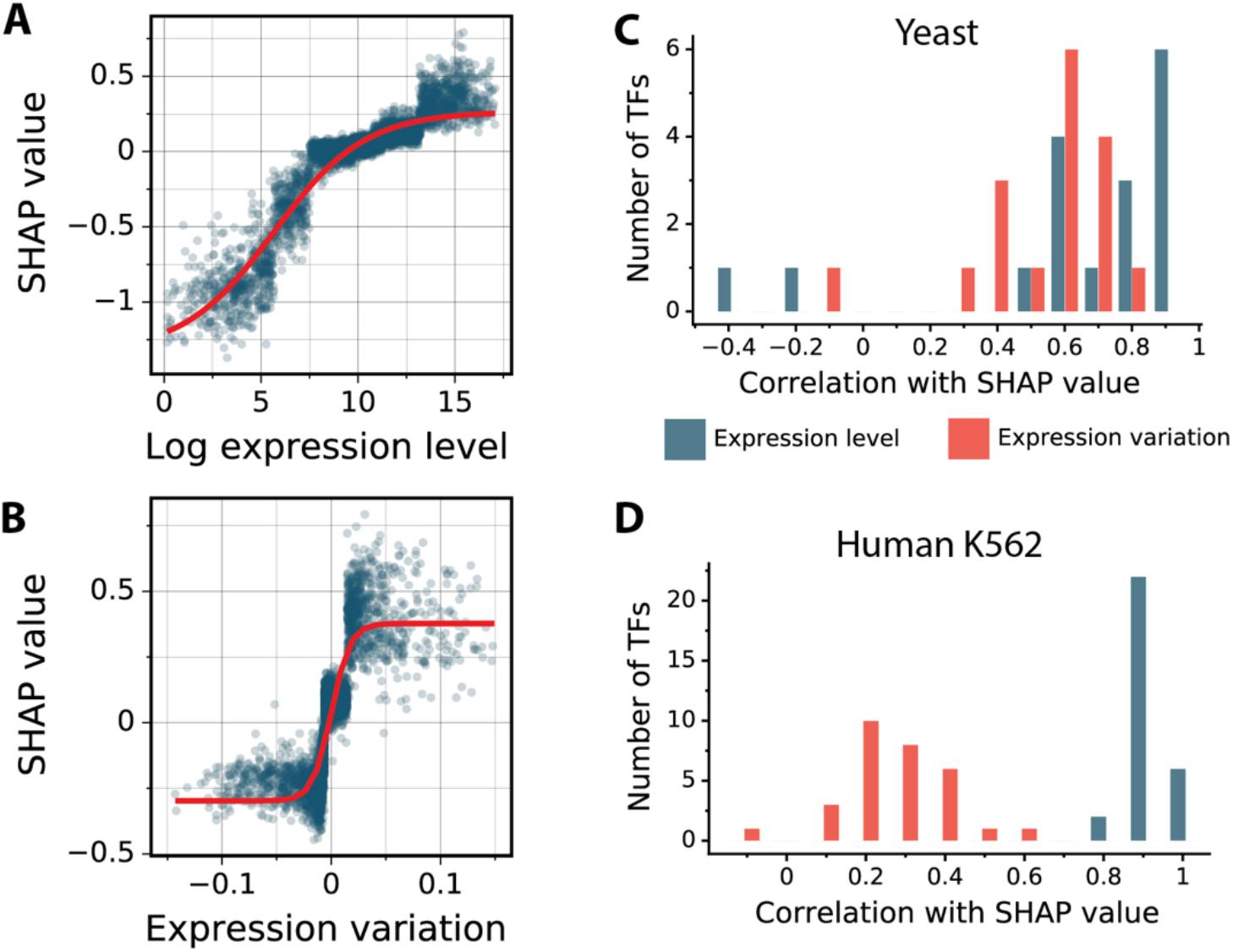
Gene-specific features. (A) Relationship between feature input and SHAP values of gene expression level for Lys14 model. Red curve is a fitted sigmoid function. The model predicts that more highly expressed genes are more likely to be responsive to Lys14 perturbation. (B) Relationship between feature input and SHAP values of gene expression variation for the same model. The model predicts that genes whose expression levels are more variable, after correction for their average expression level, are more likely to be responsive to Lys14 perturbation. (C) The distribution of the correlations between input and SHAP values for the two expression-related features in yeast cells. For most TFs, both expression level and expression variation are positively correlated with response to a perturbation. On average, expression level is more positively correlated than expression variation. All correlations are statistically significant. (D) Same as (C) for human K562 cells. All correlations are statistically significant.

### Histone marks downstream of the TSS are more predictive of responsiveness than upstream histone marks

We showed above that for human TFs, models trained using coordinate-dependent features in enhancers, the 2 kb upstream of the 5’-end TSS, and the 2 kb downstream of the 5’ TSS were no more accurate than those that used only the downstream features (Fig. 2H). For both yeast and human, the downstream histone marks had a much greater influence on the predictions than the upstream marks (Fig. 5A). This was quantified for each TF by the mean absolute SHAP values across all genes. Among the six histone marks we analyzed in yeast, downstream H3K79me1 had the biggest influence on predictions, followed by downstream H3K4me3 and downstream H3K4me1. For these three marks, the differences between their importance when they occur downstream of the TSS compared to upstream of the TSS are statistically significant (Fig. 5A, left). For human cells we did not have data on H3K79me1. Downstream H3K4me3 and H3K4me1 are the two most influential marks, followed by downstream H3K27ac (Fig. 5A, right). The differences between the importance of these marks when they occur downstream of the TSS compared to upstream are statistically significant.

**Figure 5.**
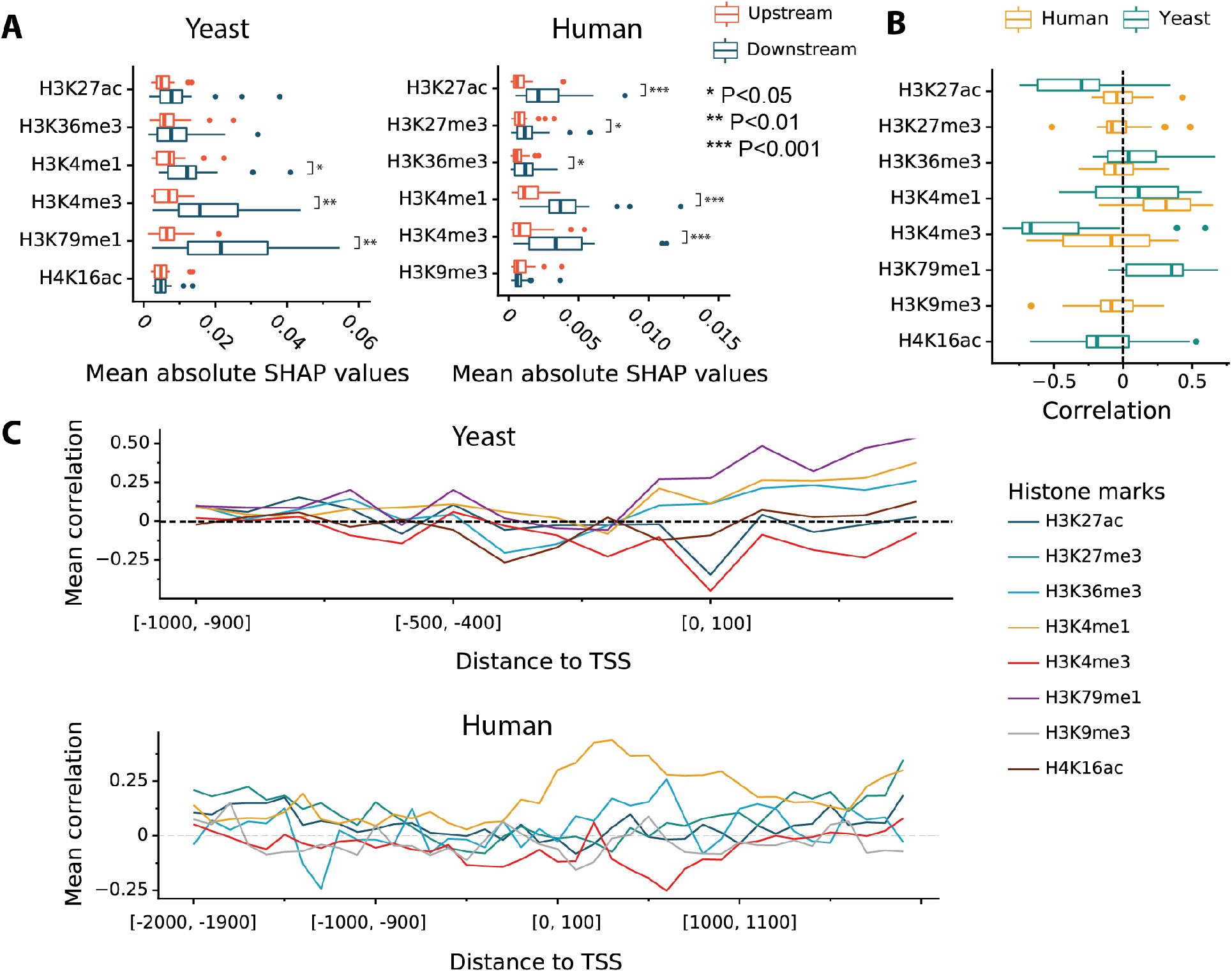
Epigenetic features. (A) Comparison of the global importance of histone mark features in the regions upstream or downstream of the TSS. The distribution across TFs is shown. For each TF, the global importance of each feature is the absolute SHAP value for that feature averaged across all genes and all promoter bins upstream or downstream of the TSS. Marks showing a significant difference are more influential when they occur downstream of the TSS than when they occur upstream. * *P*<0.05, ** *P*<0.01, *** *P*<0.001, Wilcoxon signed-rank test (B) Correlation of histone modification signals and their corresponding SHAP values at bin [0, 100] (first 100 bp downstream of the TSS). Both the yeast and human models predict that genes with stronger H3K4me1 signal immediately downstream of the TSS are more likely to be responsive to TF perturbations. Genes with stronger H3K4me3 signal, by contrast, are less likely to be responsive. (C) Average input-SHAP correlations along the genomic coordinates in yeast and human cells. In both organisms, H3K4me1 was the strongest and most consistent predictor of responsiveness while H3K4me3 was the strongest predictor of unresponsiveness.

Focusing in on the direction of influence in the 100 bp downstream of the TSS, H3K4me3 signal was negatively correlated with SHAP value for most yeast TFs, indicating that the presence of this mark pushes the model to predict a lower probability of response (Fig. 5B). H3K27ac was also negatively correlated with SHAP value for most yeast TFs, while H3K79me1 was positively correlated. For H3K4me3 and H3K4me1, the sign of correlation was generally consistent between yeast and human, but some TFs are exceptions to this generalization. Looking at different positions relative to the TSS in yeast (Fig. 5C, top), we see that the influences of H3K4me3 and H3K27ac presence are most consistently negative when they occur immediately downstream of the TSS, whereas the influences of H3K79me1, H3K4me1, and H3K36me3 presence downstream of the TSS are consistently positive. In human, the influences of H3K4m1, H3K4me3, and H3K36m3 peak slightly downstream of the TSS (Fig. 5C, bottom). However, one should not read too much into this, as the inclusion of these features in the model has a very small effect on prediction accuracy.

### Responses to any genetic perturbation are partially explained by TF-independent factors

Above, we reported that TF binding signals at cis-regulatory regions have little value for modeling responses to TF perturbations in ENCODE data on human K562 cells. This drove us to investigate whether the features that are independent of any particular TF can predict a gene’s predisposition to respond to perturbations of TFs or other regulators. We therefore calculated the response frequency of each gene – the number of perturbations to which each gene responds divided by the total number of perturbations. Next, we trained an XGBoost regression model to predict each gene’s response frequency using only the TF-independent features and tested it by 10-fold cross-validation on genes. The median variance explained is 45% for yeast and 37% for human (Fig. 6A). Since training on this task does not require DNA binding location data, we also tried it on a larger set of regulator perturbations in K562 cells, including perturbations of TFs for which no binding data is available and of regulators that are not DNA-binding proteins. This model was even better, explaining 56% of variance in the frequency of response to regulator perturbations. These results clearly demonstrate that some genes are poised to respond to perturbations and others are resistant. In this task, H3K79me1 and H3K4me3 are the most influential features in yeast cells – more influential even than gene expression level and variation (Fig. 6B). In human cells, gene expression level and variation are by far the most influential features (Fig. 6C,D). These expression features are likely read-outs of some as-yet-unidentified molecular features that make genes sensitive to perturbations.

**Figure 6.**
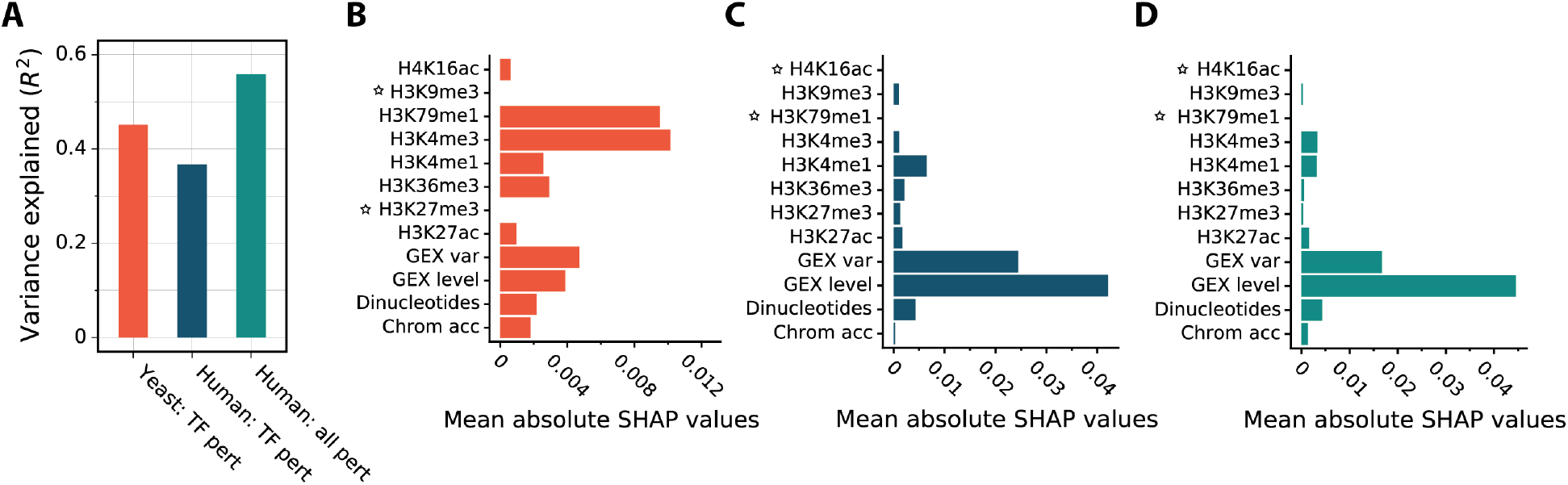
TF-independent prediction of each gene’s tendency to respond to genetic perturbations. (A) The variance explained for predicting response frequency in yeast TF perturbations (n=194), human TF perturbations for which binding data are available (n=56), and all ENCODE genetic perturbations of K562 cells (n=355). Bar height indicates the median, across genes, of the variance explained (R^2^) based on cross validation using held-out genes. (B) The mean absolute influence of each TF-independent feature on predicted response frequency, across all yeast genes. Asterisks indicate missing data. (C) Same as B for TF perturbations in human cells. (D) Same as B for all genetic perturbations in human cells.

## DISCUSSION

Determining which genes are regulated by each TF in an organism is a fundamental goal of regulatory systems biology. Furthermore, the ability to predict which genes will respond to perturbation of a TF serves as a benchmark of how well we understand the TF network. There is a body of work focused on predicting gene expression level (Middendorf et al. 2004; Ouyang et al. 2009; Schmidt et al. 2017; Tasaki et al. 2020; Karlić et al. 2010; Cheng et al. 2011; Dong et al. 2012; McLeay et al. 2012; Singh et al. 2016; Read et al. 2019; Kelley et al. 2018; Zhou et al. 2018; Washburn et al. 2019; Agarwal and Shendure 2020; Zhou et al. 2014; González et al. 2015; Crow et al. 2019; Sigalova et al. 2020), but this is a very different task from predicting the response of expression level to perturbations by using only data from unperturbed cells.

Data on where in the genome each TF binds were expected to be of great value in determining its targets, but multiple studies have shown that, in the available large ChIP-chip and ChIP-seq datasets, the genes in whose regulatory DNA a TF binds do not correspond well to those that respond to perturbation of the TF (Gitter et al. 2009; Lenstra and Holstege 2012; Cusanovich et al. 2014; Kang et al. 2020). We followed up on these observations by training machine learning models to predict which genes would respond to perturbation of a TF, given data on the TF’s binding locations and several features reflecting the gene’s epigenetic context. We found that data on yeast TF binding locations obtained by the calling cards method (Wang et al. 2011; Shively et al. 2019; Kang et al. 2020) and, to a lesser extent, the ChIP-exo method (Bergenholm et al. 2018; Holland et al. 2019), are useful for predicting which genes will respond to a perturbation of the TF. In fact, binding location was the most influential and valuable among the features we provided (Fig. 2A-D). Since earlier ChIP-chip data on yeast are known to correspond poorly to perturbation response, we conclude that the newer technologies are yielding better results. Binding signals influenced predictions mainly in the 500 bp upstream of the TSS, suggesting that this is the extent of functional yeast promoter regions (Fig. 3A-B). Even among genes with significant binding signal for a TF in their promoter, the strength and location of the signal helped to differentiate between functional and non-functional binding (Fig 3C-D). In ENCODE data on human K562 cells, however, the situation was strikingly different. The models did not identify any patterns in ChIP-seq data on the perturbed TF that were useful for predicting which genes would respond to the perturbation (Fig. 2E-F).

We also investigated the predictive value of selected histone marks and chromatin accessibility features. In yeast, these features had predictive value that was less than that of TF binding locations, but was not negligible (Fig. 2A-D). In human, however, these features were not useful in models that included pre-perturbation gene expression level and variation (GEX features). HM features had a larger (though still small) impact in models that did not include GEX features (discussed below). Among HMs for which we had data in both yeast and human, H3K4me1 and H3K4me3 were most influential. H3K4me1 increased the predicted probability of a gene’s response to perturbations while H3K4me3 decreased it (Fig. 5). Both marks were most influential when they occurred downstream of the TSS, in the gene body. In fact, dropping all features that mapped to the enhancers and the promoter region upstream of the TSS had only a small impact on predictive accuracy in K562 cells. This surprising observation likely reflects both the low utility of the existing ChIP-seq data and incomplete knowledge of enhancer locations and enhancer-gene associations. Future datasets on TF binding locations and enhancer-gene associations will likely reveal at least some predictive power for enhancer features.

Pre-perturbation gene expression level and variation were surprisingly good predictors of which genes would respond to a perturbation. In fact, a model using only these two features predicted responses in K562 cells almost as well as the full model, which includes TF ChIP-seq, histone marks, chromatin accessibility, and dinucleotide frequencies (Fig. 2F). Genes that were expressed at a higher level and genes that showed more variability in their expression level were more likely to respond to perturbations (Fig. 4). We hypothesize that these features are readouts of molecular features of each gene’s sequence context and / or epigenetic state which have limited predictive power individually, but much greater predictive power when aggregated by their effects on gene expression level and variation. This hypothesis is supported by the observation that the influence of several histone marks increases when GEX features are omitted from the model, though these influences are still small compared to the impact of GEX features (Fig. 2G). However, the epigenetic state that is reflected in the GEX features is not as simple open chromatin versus closed chromatin, since chromatin accessibility has little influence even in the absence of GEX features. Previous studies suggest that several sequence features, including the presence of a TATA box in the promoter (Blake et al. 2006; Ravarani et al. 2016; Sigalova et al. 2020) and the GC content of the promoter (Morgan and Marioni 2018; Sigalova et al. 2020) influence expression variation. Identifying additional sequence elements or epigenetic states that are reflected in GEX features and showing that they can predict perturbation response is an important direction for future research.

The predictive power of features that are independent of the TF perturbed -- GEX features, HMs, chromatin accessibility, and dinucleotide sequence -- shows that some genes are poised to respond to perturbations while others are not. To confirm this, we trained models to predict how many TF perturbations a gene would respond to by using only these features of the target gene (not TF binding locations). This model proved quite accurate (Fig. 6A). The yeast model relied most on histone marks, followed by GEX features (Fig. 6B), whereas the human model relied almost exclusively on GEX features (Fig. 6C-D). This was a significant finding -- if you want to predict which genes will respond to perturbation of a TF, it is not enough to know where the TF binds -- you must also know whether the gene is predisposed to respond to perturbations. An important direction for future research is to discover the extent to which a gene’s predisposition to respond to perturbations is an epigenetic feature that depends on cell type and conditions or an inherent feature of the gene.

Neither the yeast nor the human model is able to predict which genes will respond to a TF perturbation reliably, even when provided with the binding locations of the perturbed TF and a host of epigenetic features. This is not simply a consequence of discretizing perturbation responsiveness, as models trained to predict the quantitative response yielded similar results (not shown). One possible route to improving accuracy would be to provide binding location features for other TFs that interact in some way with the perturbed TF. For example, some genes may respond to perturbation of TF that does not bind in their regulatory DNA because the perturbed TF regulates another TF that does bind in their regulatory DNA. In principle, a learning algorithm given the binding locations of all TFs might be able to discover such network effects itself, but in practice the number of features is likely to be too large compared to the number of training examples for responses to perturbation of a single TF. An alternative would be to map the TF network externally, using NetProphet 2, Inferelator, or other network inference algorithms (Kang et al. 2018; Greenfield et al. 2013; Roy et al. 2013; Huynh-Thu et al. 2010). The response prediction algorithm could then be provided with binding location data on TFs that the perturbed TF regulates. With those data, it could learn that genes respond to perturbation of a given TF if they are bound by another TF. Expanding to other types of TF-TF interactions, binding data might also be for TFs that interact physically with the perturbed TF, according to protein-protein interaction maps (Szklarczyk et al. 2019; Oughtred et al. 2019).

For human cells, the inability to predict responsiveness may also reflect limitations of the technologies used for measuring TF binding locations and perturbing TFs, as well as limited knowledge of enhancer locations and enhancer-gene associations. It may also be possible to improve on the way enhancer-associated features were coded, enabling models to better utilize histone marks and chromatin accessibility for determining which enhancers are active in a given sample of cells (Fullwood and Ruan 2009; Mumbach et al. 2016, 2017; Fulco et al. 2016; Klann et al. 2017; Simeonov et al. 2017; Lonsdale et al. 2013; Aguet et al. 2017). Other types of data, such as levels of enhancer-associated transcription, may also help (Core et al. 2014; Mahat et al. 2016; Tome et al. 2018; Azofeifa et al. 2018). For yeast, however, these explanations are less applicable. Obtaining the right genomic data and developing the right models for predicting which yeast genes will respond to perturbation of a TF remains a major challenge. Progress in overcoming this challenge will serve as a benchmark for our understanding of regulatory systems biology.

## METHODS

Code is available at https://github.com/yiming-kang/TFPertRespExplainer. All data and code will be archived on a Zenodo page that is under construction. Additional details are provided in Supplemental Methods.

### TF-perturbation response data

We downloaded yeast microarray data taken 15 minutes after inducing overexpression of 194 TFs from https://idea.research.calicolabs.com/data (Hackett et al. 2020). Column *log2_shrunken_timecourses* from file “Raw & processed gene expression data” was used. Since these values were already shrunken towards zero, any gene with a non-zero value was defined as responsive. For human K562 cells, we used all RNA-seq expression profiles after gene perturbation in K562 cells from the ENCODE Project database (Dunham et al. 2012; Davis et al. 2018; Abascal et al. 2020). Perturbations include disabling mutations introduced by CRISPR, CRISPR inference, small-interfering RNA, and small-hairpin RNA. We downloaded RSEM expected counts of experimental and control profiles. For each of the 355 experiments, we ran DESeq2 (V1.10.1) (Love et al. 2014) to identify differentially expressed genes by comparing the experimental replicates to the corresponding control replicates. Genes with adjusted *P* < 0.05 and absolute log2 fold-change > 0.5 were considered responsive.

### Pre-perturbation gene expression features

The pre-perturbation expression level feature of a gene is its median expression level across all samples measured prior to the TF perturbation. For human genes (RNA-Seq data) we averaged the log TPM levels of all control replicates. For yeast (microarray data) we used log fluorescence levels of the red (experimental) channel measured at time 0 (before TF induction). To construct a gene expression variation feature that is independent of the expression level, we used the method of (Sigalova et al. 2020).

### TF binding location features

All coordinate-dependent features were mapped to yeast genome build sacCer3 and human build GRCh38. Yeast binding location data were generated using transposon calling cards (16 TFs) (Wang et al. 2011; Shively et al. 2019; Kang et al. 2020) or ChIP-exo (20 TFs) (Bergenholm et al. 2018; Holland et al. 2019). ChIP-exo were obtained directly from the authors of ref. (Bergenholm et al. 2018; Holland et al. 2019). Following ref. (Kang et al. 2020), we focused on the ChIP-exo data from glucose limited chemostats. Peak locations reported for strain CEN.PK were transferred to SacCer3 coordinates (strain S288C) as described in Supplemental Methods. We downloaded ChIP-seq peaks for human 54 TFs in K562 cells from ENCODE. K562 had far more TFs that were both ChIPped and perturbed than any other cell type. We consider a gene to be a TF only if it has a well-defined DNA-binding domain (Lambert et al. 2018). We used the “conservative” peaks, which have Irreproducible Discovery Rate <= 2%. The log10 q-value reported for each peak was used as the binding signal feature.

### Histone modifications and chromatin accessibility data

For yeast, we used histone modification data at timepoint 0 (in YPD batch cultures prior to addition of a diamide stress) from ref. (Weiner et al. 2015), which was produced using MNase-ChIP-Seq (GEO accession GSE61888, https://www.ncbi.nlm.nih.gov/geo/). We used chromatin accessibility data at timepoint 0 (prior to addition of osmotic stress) from ref. (Schep et al. 2015) (GSE66386). For human K562 cells, we downloaded the coverage data (fold change over control) for histone modifications and chromatin accessibility from ENCODE.

### Mapping genome-wide features to cis-regulatory regions

We defined yeast promoter regions as 1,000 bp upstream to 500 bp downstream from the transcription start site (TSS). TSS coordinates were obtained from ref. (de Boer et al. 2020). Inputs for each genome-wide feature were mapped relative to the TSS and summed within each of 15 100 bp bins to create 15 features. Human TSS coordinates were downloaded from Ensembl Release 92 (Cunningham et al. 2019). For each gene, we defined the *5’ promoter* to be 4 Kb centered on the 5’-end TSS. We define *alternative promoters* to be 4 Kb regions centered at any TSS that is more than 2 Kb from the 5’-end TSS and *enhancers* to be enhancers that are linked to the gene in GeneHancer V4.8 (Fishilevich et al. 2017). We used only “double elite” enhancers, whose existence and linkage to the target gene are both supported by at least two types of evidence.

Each gene must have an equal number of features to create a rectangular feature matrix, even though genes differ in how many alternative TSSs and enhancers they have. For TSSs, we primarily focused on 5’-end (see Supplemental Methods for rationale). Alternative TSSs within 2 Kb of the 5’ end were deemed to be included in the 5’ promoter and those further than 2 Kb were treated as enhancers, since enhancers and promoters share most properties and functions (Andersson and Sandelin 2020). Signals located in enhancers were aggregated into features in two ways. The first method (*Prom + bin enhan*; Fig. S3, blue) sums signals within each of 45 bins upstream of the 5’ end promoter region and another 45 bins downstream. The width of the bin closest to the TSS was 500 bp and each subsequent bin was larger by 500 bp (500 bp, 1 Kb, 1.5 Kb, etc.). Together with the 5’ promoter, these bins 500 Kb on either side of the 5’ TSS. The second method (*Prom + agg enhan*, Fig. S3, green) sums enhancer signals 500 Kb upstream or downstream of the promoter into two features. Only signals within defined enhancers were used.

### Training and Testing Machine Learning Algorithms

For each TF perturbation, we trained and tested a model for predicting whether a gene will respond by using 10-fold cross-validation. The genes in each fold were selected at random subject to the constraint that all folds have the same proportion of responsive genes. We tried two ensemble classification algorithms—random forests implemented in scikit-learn library (V0.22.1) (Pedregosa et al. 2011), and gradient boosted trees implemented in XGBoost library (V 0.90) (Chen and Guestrin 2016). For both methods, 500 trees were used. For XGBoost, learning rate of 0.01 was used for the “gbtree” booster. All other parameters were default. See Supplemental Methods for details.

### Using SHAP to quantify the influences of features on predictions

We employed the SHAP (SHapley Additive exPlanations) framework (V0.35.0) (Lundberg and Lee 2017) to quantify the influence of each feature on each prediction (TreeExplainer function). SHAP uses a linear model to approximate the predictions for artificially constructed instances in the neighborhood of each actual instance. SHAP values explain why one particular prediction – one TF-gene pair – differs from the average prediction for all genes in response to perturbation of that TF. To characterize the effect of a particular feature for a set of genes, we separately summed its positive and negative SHAP values over genomic bins for each gene. We then averaged the positive sums over all genes and, separately, the negative sums.

### Modeling and interpreting generic responses in any genetic perturbation

We used an XGBoost regressor to predict the fraction of genetic perturbations to which each gene would respond, in absence of perturbation-specific information. Three data sets consisted of all TF induction samples in yeast (n=194), all TF perturbation samples in K562 cells (n=56), and all genetic perturbations in K562 (n=355). Performance was evaluated by cross validation as described above.

## COMPETING INTERESTS

The authors declare that they have no competing interests.

## AVAILABILITY OF DATA AND MATERIALS

Code is available at https://github.com/yiming-kang/TFPertRespExplainer. All data were downloaded from the publicly available sources indicated in Methods. All data, code, and analyses will be archived on a Zenodo page that is under construction.

## Notes

### Competing Interest Statement

The authors have declared no competing interest.

## REFERENCES

Abascal F, Acosta R, Addleman NJ, Adrian J, Afzal V, Aken B, Akiyama JA, Jammal O Al, Amrhein H, Anderson SM, et al. 2020. Expanded encyclopaedias of DNA elements in the human and mouse genomes. Nature 583: 699–710.

Agarwal V, Shendure J. 2020. Predicting mRNA Abundance Directly from Genomic Sequence Using Deep Convolutional Neural Networks. Cell Rep 31: 107663. https://doi.org/10.1016/j.celrep.2020.107663.

Aguet F, Brown AA, Castel SE, Davis JR, He Y, Jo B, Mohammadi P, Park YS, Parsana P, Segrè A V., et al. 2017. Genetic effects on gene expression across human tissues. Nature.

Andersson R, Sandelin A. 2020. Determinants of enhancer and promoter activities of regulatory elements. Nat Rev Genet 21: 71–87. http://dx.doi.org/10.1038/s41576-019-0173-8.

Azofeifa JG, Allen MA, Hendrix JR, Read T, Rubin JD, Dowell RD. 2018. Enhancer RNA profiling predicts transcription factor activity. Genome Res 28: 334–344.

Bergenholm D, Liu G, Holland P, Nielsen J. 2018. Reconstruction of a Global Transcriptional Regulatory Network for Control of Lipid Metabolism in Yeast by Using Chromatin Immunoprecipitation with Lambda Exonuclease Digestion. mSystems.

Breiman L. 2001. Random forests. Mach Learn.

Chen T, Guestrin C. 2016. XGBoost: A scalable tree boosting system. In Proceedings of the ACM SIGKDD International Conference on Knowledge Discovery and Data Mining.

Cheng C, Yan KK, Yip KY, Rozowsky J, Alexander R, Shou C, Gerstein M. 2011. A statistical framework for modeling gene expression using chromatin features and application to modENCODE datasets. Genome Biol.

Core LJ, Martins AL, Danko CG, Waters CT, Siepel A, Lis JT. 2014. Analysis of nascent RNA identifies a unified architecture of initiation regions at mammalian promoters and enhancers. Nat Genet.

Crow M, Lim N, Ballouz S, Pavlidis P, Gillis J. 2019. Predictability of human differential gene expression. Proc Natl Acad Sci U S A 116: 6491–6500.

Cunningham F, Achuthan P, Akanni W, Allen J, Amode MR, Armean IM, Bennett R, Bhai J, Billis K, Boddu S, et al. 2019. Ensembl 2019. Nucleic Acids Res.

Cusanovich DA, Pavlovic B, Pritchard JK, Gilad Y. 2014. The Functional Consequences of Variation in Transcription Factor Binding. PLoS Genet 10.

Davis CA, Hitz BC, Sloan CA, Chan ET, Davidson JM, Gabdank I, Hilton JA, Jain K, Baymuradov UK, Narayanan AK, et al. 2018. The Encyclopedia of DNA elements (ENCODE): Data portal update. Nucleic Acids Res.

de Boer CG, Vaishnav ED, Sadeh R, Abeyta EL, Friedman N, Regev A. 2020. Deciphering eukaryotic gene-regulatory logic with 100 million random promoters. Nat Biotechnol 38: 56–65. http://dx.doi.org/10.1038/s41587-019-0315-8.

Dong X, Greven MC, Kundaje A, Djebali S, Brown JB, Cheng C, Gingeras TR, Gerstein M, Guigó R, Birney E, et al. 2012. Modeling gene expression using chromatin features in various cellular contexts. Genome Biol.

Dunham I, Kundaje A, Aldred SF, Collins PJ, Davis CA, Doyle F, Epstein CB, Frietze S, Harrow J, Kaul R, et al. 2012. An integrated encyclopedia of DNA elements in the human genome. Nature.

Fisher A, Rudin C, Dominici F. 2019. All models are wrong, but many are useful: Learning a variable’s importance by studying an entire class of prediction models simultaneously. J Mach Learn Res.

Fishilevich S, Nudel R, Rappaport N, Hadar R, Plaschkes I, Iny Stein T, Rosen N, Kohn A, Twik M, Safran M, et al. 2017. GeneHancer: genome-wide integration of enhancers and target genes in GeneCards. Database 2017: 1–17. https://academic.oup.com/database/article-lookup/doi/10.1093/database/bax028.

Fulco CP, Munschauer M, Anyoha R, Munson G, Grossman SR, Perez EM, Kane M, Cleary B, Lander ES, Engreitz JM. 2016. Systematic mapping of functional enhancer-promoter connections with CRISPR interference. Science (80-).

Fullwood MJ, Ruan Y. 2009. ChIP-based methods for the identification of long-range chromatin interactions. J Cell Biochem.

Gitter A, Siegfried Z, Klutstein M, Fornes O, Oliva B, Simon I, Bar-Joseph Z. 2009. Backup in gene regulatory networks explains differences between binding and knockout results. Mol Syst Biol 5: 1–7. http://dx.doi.org/10.1038/msb.2009.33.

González AJ, Setty M, Leslie CS. 2015. Early enhancer establishment and regulatory locus complexity shape transcriptional programs in hematopoietic differentiation. Nat Genet.

Grant CE, Bailey TL, Noble WS. 2011. FIMO: Scanning for occurrences of a given motif. Bioinformatics 27: 1017–1018.

Greenfield A, Hafemeister C, Bonneau R. 2013. Robust data-driven incorporation of prior knowledge into the inference of dynamic regulatory networks. Bioinformatics 29: 1060–1067.

Hackett SR, Baltz EA, Coram M, Wranik BJ, Kim G, Baker A, Fan M, Hendrickson DG, Berndl M, Mcisaac RS. 2020. Learning causal networks using inducible transcription factors and transcriptome-wide time series. 1–15.

Henikoff S, Shilatifard A. 2011. Histone modification: Cause or cog? Trends Genet 27: 389–396. http://dx.doi.org/10.1016/j.tig.2011.06.006.

Holland P, Bergenholm D, Börlin CS, Liu G, Nielsen J. 2019. Predictive models of eukaryotic transcriptional regulation reveals changes in transcription factor roles and promoter usage between metabolic conditions. Nucleic Acids Res.

Huynh-Thu VA, Irrthum A, Wehenkel L, Geurts P. 2010. Inferring regulatory networks from expression data using tree-based methods. PLoS One.

Japkowicz N, Stephen S. 2002. The class imbalance problem: A systematic study. Intell Data Anal.

Kang Y, Liow HH, Maier EJ, Brent MR. 2018. NetProphet 2.0: Mapping transcription factor networks by exploiting scalable data resources. Bioinformatics 34: 249–257.

Kang Y, Patel NR, Shively C, Recio PS, Chen X, Wranik BJ, Kim G, McIsaac RS, Mitra R, Brent MR. 2020. Dual threshold optimization and network inference reveal convergent evidence from TF binding locations and TF perturbation responses. Genome Res gr.259655.119.

Karlić R, Chung HR, Lasserre J, Vlahoviček K, Vingron M. 2010. Histone modification levels are predictive for gene expression. Proc Natl Acad Sci U S A 107: 2926–2931.

Kelley DR, Reshef YA, Bileschi M, Belanger D, Mclean CY, Snoek J. 2018. Sequential regulatory activity prediction across chromosomes with convolutional neural networks. 1–12.

Klann TS, Black JB, Chellappan M, Safi A, Song L, Hilton IB, Crawford GE, Reddy TE, Gersbach CA. 2017. CRISPR-Cas9 epigenome editing enables high-throughput screening for functional regulatory elements in the human genome. Nat Biotechnol.

Lambert SA, Jolma A, Campitelli LF, Das PK, Yin Y, Albu M, Chen X, Taipale J, Hughes TR, Weirauch MT. 2018. The Human Transcription Factors. Cell 172: 650–665. https://doi.org/10.1016/j.cell.2018.01.029.

Lenstra TL, Holstege FCP. 2012. The discrepancy between chromatin factor location and effect. Nucl (United States).

Lonsdale J, Thomas J, Salvatore M, Phillips R, Lo E, Shad S, Hasz R, Walters G, Garcia F, Young N, et al. 2013. The Genotype-Tissue Expression (GTEx) project. Nat Genet.

Love MI, Huber W, Anders S. 2014. Moderated estimation of fold change and dispersion for RNA-seq data with DESeq2. Genome Biol 15: 1–21.

Lundberg S, Lee S-I. 2017. A Unified Approach to Interpreting Model Predictions. NIPS 16: 426–430.

Lundberg SM, Erion GG, Lee S. 2018. Consistent Individualized Feature Attribution for Tree Ensembles. http://arxiv.org/abs/1802.03888.

Mahat DB, Kwak H, Booth GT, Jonkers IH, Danko CG, Patel RK, Waters CT, Munson K, Core LJ, Lis JT. 2016. Base-pair-resolution genome-wide mapping of active RNA polymerases using precision nuclear run-on (PRO-seq). Nat Protoc.

McLeay RC, Lesluyes T, Cuellar Partida G, Bailey TL. 2012. Genome-wide in silico prediction of gene expression. Bioinformatics.

Middendorf M, Kundaje A, Wiggins C, Freund Y, Leslie C. 2004. Predicting genetic regulatory response using classification. In Bioinformatics.

Molnar C. 2019. Interpretable Machine Learning. A Guide for Making Black Box Models Explainable. Book.

Mumbach MR, Rubin AJ, Flynn RA, Dai C, Khavari PA, Greenleaf WJ, Chang HY. 2016. HiChIP: efficient and sensitive analysis of protein-directed genome architecture. Nat Methods 13: 919–922. http://www.ncbi.nlm.nih.gov/pubmed/27643841%0A http://www.pubmedcentral.nih.gov/articlerender.fcgi?artid=PMC5501173.

Mumbach MR, Satpathy AT, Boyle EA, Dai C, Gowen BG, Cho SW, Nguyen ML, Rubin AJ, Granja JM, Kazane KR, et al. 2017. Enhancer connectome in primary human cells identifies target genes of disease-associated DNA elements. Nat Genet 49: 1602–1612. http://www.ncbi.nlm.nih.gov/pubmed/28945252%0A http://www.pubmedcentral.nih.gov/articlerender.fcgi?artid=PMC5805393.

Oughtred R, Stark C, Breitkreutz BJ, Rust J, Boucher L, Chang C, Kolas N, O’Donnell L, Leung G, McAdam R, et al. 2019. The BioGRID interaction database: 2019 update. Nucleic Acids Res.

Ouyang Z, Zhou Q, Wong WH. 2009. ChIP-Seq of transcription factors predicts absolute and differential gene expression in embryonic stem cells. Proc Natl Acad Sci U S A.

Pedregosa F, Varoquaux G, Gramfort A, Michel V, Thirion B, Grisel O, Blondel M, Prettenhofer P, Weiss R, Dubourg V, et al. 2011. Scikit-learn: Machine learning in Python. J Mach Learn Res.

Read DF, Cook K, Lu YY, Le Roch KG, Noble WS. 2019. Predicting gene expression in the human malaria parasite Plasmodium falciparum using histone modification, nucleosome positioning, and 3D localization features. PLoS Comput Biol 15: 1–23. http://dx.doi.org/10.1371/journal.pcbi.1007329.

Roadmap Epigenomics Consortium, Kundaje A, Meuleman W, Ernst J, Bilenky M, Yen A, Heravi-Moussavi A, Kheradpour P, Zhang Z, Wang J, et al. 2015. Integrative analysis of 111 reference human epigenomes. Nature.

Rossi MJ, Lai WKM, Pugh BF. 2018. Genome-wide determinants of sequence-specific DNA binding of general regulatory factors. Genome Res.

Roy S, Lagree S, Hou Z, Thomson JA, Stewart R, Gasch AP. 2013. Integrated Module and Gene-Specific Regulatory Inference Implicates Upstream Signaling Networks. PLoS Comput Biol 9.

Schep AN, Buenrostro JD, Denny SK, Schwartz K, Sherlock G, Greenleaf WJ. 2015. Structured nucleosome fingerprints enable high-resolution mapping of chromatin architecture within regulatory regions. Genome Res.

Schmidt F, Gasparoni N, Gasparoni G, Gianmoena K, Cadenas C, Polansky JK, Ebert P, Nordstrom K, Barann M, Sinha A, et al. 2017. Combining transcription factor binding affinities with open-chromatin data for accurate gene expression prediction. Nucleic Acids Res.

Shively CA, Liu J, Chen X, Loell K, Mitra RD. 2019. Homotypic cooperativity and collective binding are determinants of bHLH specificity and function. Proc Natl Acad Sci U S A.

Sigalova O, Shaeiri A, Forneris M, Furlong E, Zaugg J. 2020. Predictive features of gene expression variation reveal a mechanistic link between expression variation and differential expression. 1–24.

Simeonov DR, Gowen BG, Boontanrart M, Roth TL, Gagnon JD, Mumbach MR, Satpathy AT, Lee Y, Bray NL, Chan AY, et al. 2017. Discovery of stimulation-responsive immune enhancers with CRISPR activation. Nature.

Singh R, Lanchantin J, Robins G, Qi Y. 2016. DeepChrome: Deep-learning for predicting gene expression from histone modifications. In Bioinformatics.

Spivak AT, Stormo GD. 2012. ScerTF: A comprehensive database of benchmarked position weight matrices for Saccharomyces species. Nucleic Acids Res.

Szklarczyk D, Gable AL, Lyon D, Junge A, Wyder S, Huerta-Cepas J, Simonovic M, Doncheva NT, Morris JH, Bork P, et al. 2019. STRING v11: Protein-protein association networks with increased coverage, supporting functional discovery in genome-wide experimental datasets. Nucleic Acids Res.

Tasaki S, Gaiteri C, Mostafavi S, Wang Y. 2020. Deep learning decodes the principles of differential gene expression. Nat Mach Intell.

Tome JM, Tippens ND, Lis JT. 2018. Single-molecule nascent RNA sequencing identifies regulatory domain architecture at promoters and enhancers. Nat Genet.

Wang H, Mayhew D, Chen X, Johnston M, Mitra RD. 2011. Calling Cards enable multiplexed identification of the genomic targets of DNA-binding proteins. Genome Res.

Washburn JD, Mejia-Guerra MK, Ramstein G, Kremling KA, Valluru R, Buckler ES, Wang H. 2019. Evolutionarily informed deep learning methods for predicting relative transcript abundance from DNA sequence. Proc Natl Acad Sci U S A 116: 5542–5549.

Weiner A, Hsieh TS, Rando OJ, Friedman N, Weiner A, Hsieh TS, Appleboim A, Chen H V, Rahat A, Amit I. 2015. High-Resolution Chromatin Dynamics during a Yeast Resource High-Resolution Chromatin Dynamics during a Yeast Stress Response. Mol Cell 58: 371–386. http://dx.doi.org/10.1016/j.molcel.2015.02.002.

Zeiler MD, Fergus R. 2012. Visualizing and Understanding Convolutional Networks.

Zhou J, Theesfeld CL, Yao K, Chen KM, Wong AK, Troyanskaya OG. 2018. Deep learning sequence-based ab initio prediction of variant effects on expression and disease risk. Nat Genet.

Zhou J, Troyanskaya OG. 2015. Predicting effects of noncoding variants with deep learning–based sequence model. Nat Methods.

Zhou X, Cain CE, Myrthil M, Lewellen N, Michelini K, Davenport ER, Stephens M, Pritchard JK, Gilad Y. 2014. Epigenetic modifications are associated with inter-species gene expression variation in primates. Genome Biol.

