## Supplemental Materials for "Predicting which genes will respond to perturbations of a TF: TF-independent properties of genes are major determinants of their responsiveness"

**SUPPLEMENTAL FIGURES**

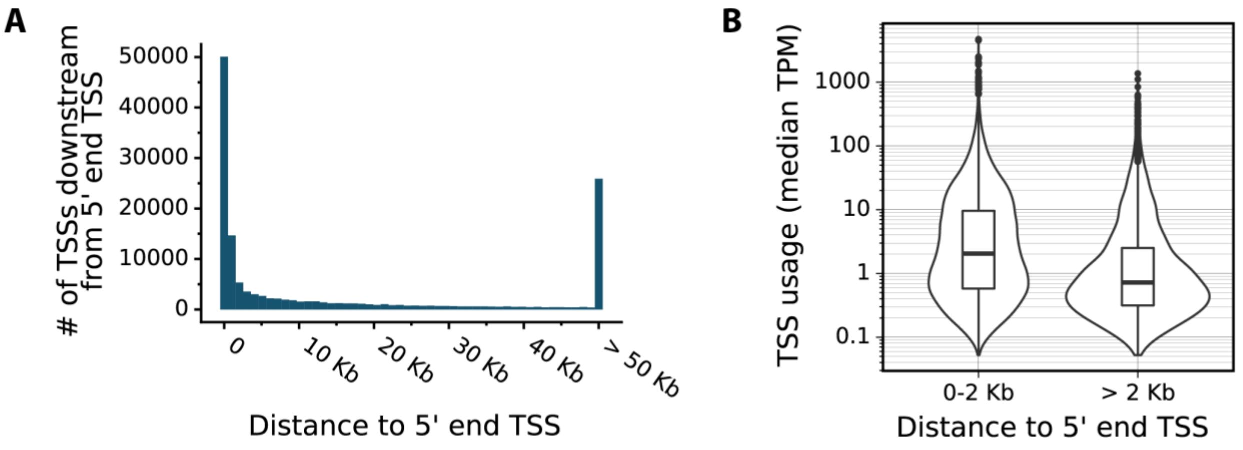


**Supplemental Figure S1.** Statistics on human TSS. (A) Distances between each 5’ end TSS and other downstream TSSs of the corresponding gene. Among the downstream TSSs for all genes, ~47% of them are within the 2 Kb range of their paired 5’ end TSS. The median distance for the TSSs within 2 Kb range is 163 bp, while the median distance for those that are more than 2 Kb away is 26.3 Kb. (B) Relationship between TSS usage and distance. TSS usage for each TSS is represented as the median Tags Per Million (TPM) level across all samples in Fantom5 CAGE expression data (Forrest et al. 2014; Lizio et al. 2019). The median TPM for the TSSs within 2 Kb range to their corresponding 5’ end TSS (including these 5’ end TSS themselves) is approximately three times of the median TPM for the TSSs that are more than 2 Kb away from their corresponding 5’ end TSS.

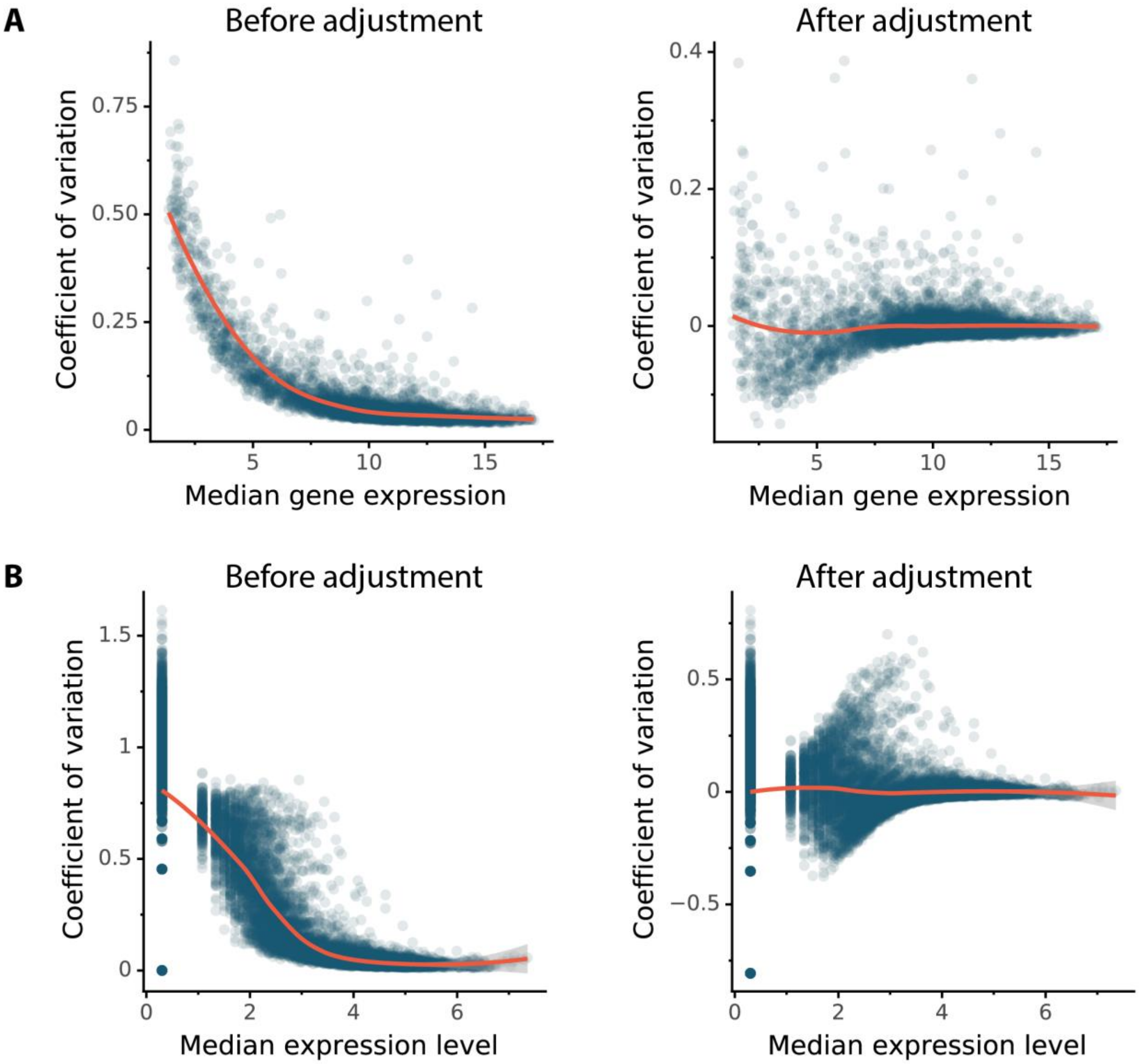


**Supplemental Figure S2.** (A) Expression variation for yeast cells. Left: Relationship between the median expression level of each gene across pre-perturbation (or control) conditions and its expression variation measured by the coefficient of variation. Orange curve was fitted using locally estimated scatterplot smoothing (LOESS) regression. Right: Expression variation adjusted for the median expression level by taking the residual of LOESS regression (the orange curve from left). (B) Expression variation for human cells. Same analysis as in (A).

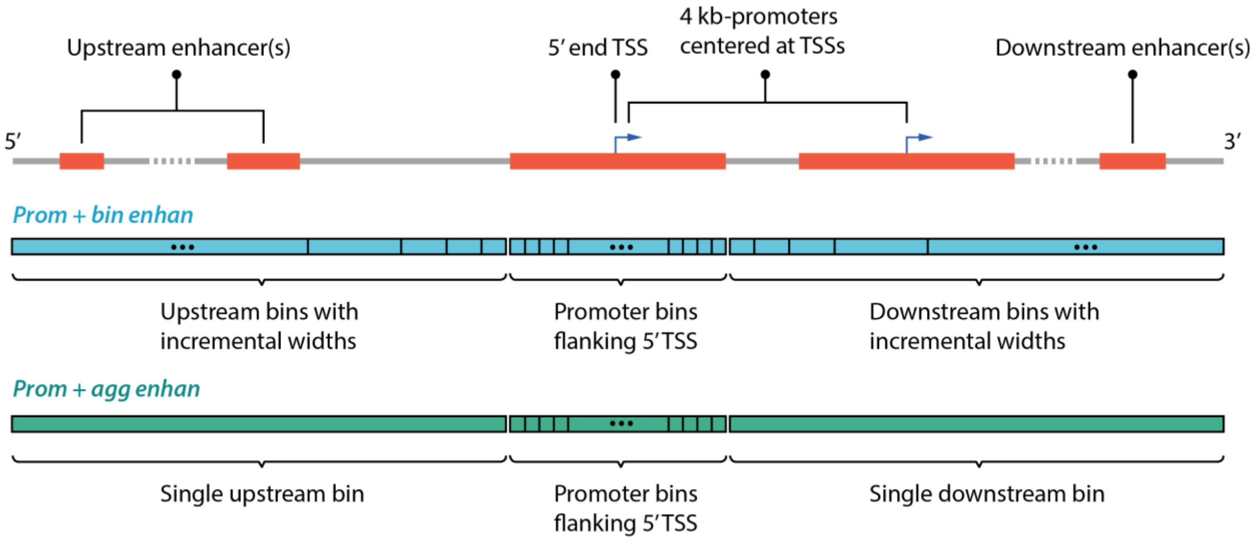


**Supplemental Figure S3.** Definition of human cis-regulatory regions. The top panel illustrates a 1 Mb region centered at the 5' end TSS of a gene. Orange boxes indicate the enhancers linked to the gene, and the 4 Kb promoter(s) centered around the gene's TSS(s). The bottom two panels illustrate the approaches for binning the 1Mb cis-regulatory region. *Prom + bin enhan* (blue) includes 40 equal-sized bins of the promoter centered around the 5' TSS, and 45 bins with incremental widths for the upstream regions between -500 Kb and -2 Kb and another 45 bins for the downstream regions between 2 Kb and 500 Kb respectively. As the distance between the distal bin and TSS increases, the width of the bin increases exponentially. *Prom + agg enhan* (green) includes 40 equal-sized bins of the promoter centered around the 5' TSS, one single upstream bin covering the entire region between -500 Kb and -2 Kb, and one single downstream bin covering the region between 2 Kb and 500 Kb. The signals of each coordinate-dependent feature that are mapped to the defined cis-regulatory regions (orange, top panel) linked to the corresponding bins according to the genomic coordinates. Within each bin, the signals are summed into a single aggregated input value.

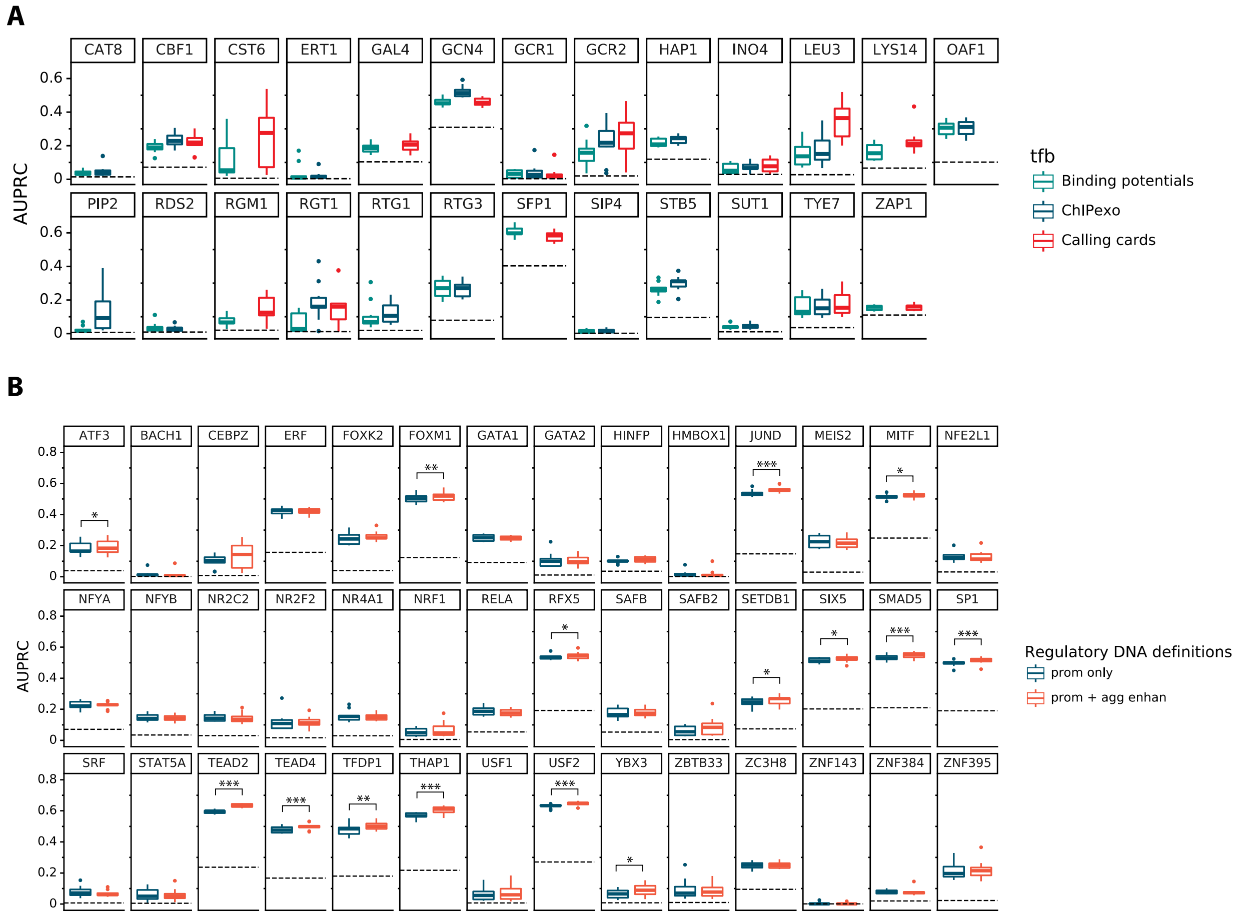


**Supplemental Figure S4.** (A) Performance of individual yeast TF models that were trained on different types of TF binding data. No boxplot is shown if the TF binding data from the corresponding assay was unavailable. Each boxplot shows the results of ten-fold cross-validation on all genes. (B) Performance of individual human TF models that were trained using various definitions of regulatory DNA. Statistical significance used paired t-test: $p < 0.05$ (*), $p < 0.01$ (**), $p < 0.001$(***).

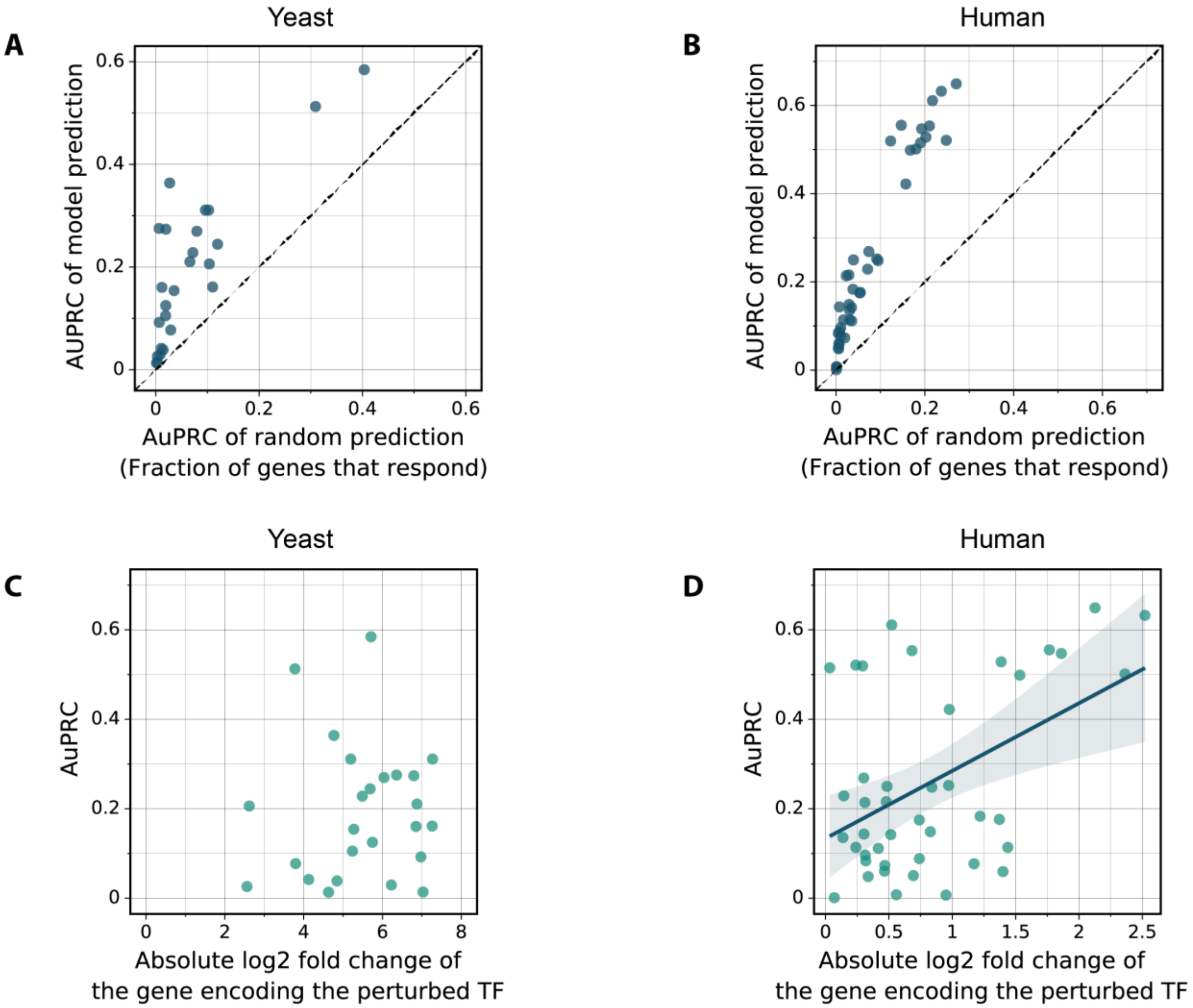


**Supplemental Figure S5.** (A) Relationship between AUPRC of random prediction and AUPRC of model prediction. Dashed diagonal line has slope of 1. (B) Same as (A) but for human K562 TFs. (C) Relationship between the log fold change of the mRNA for the perturbed TF and model accuracy, for yeast. Pearson correlation = 0.06, $P$ = 0.76. (D) Same as (B) but for human TFs. Pearson correlation = 0.47, $P$ = 0.002.

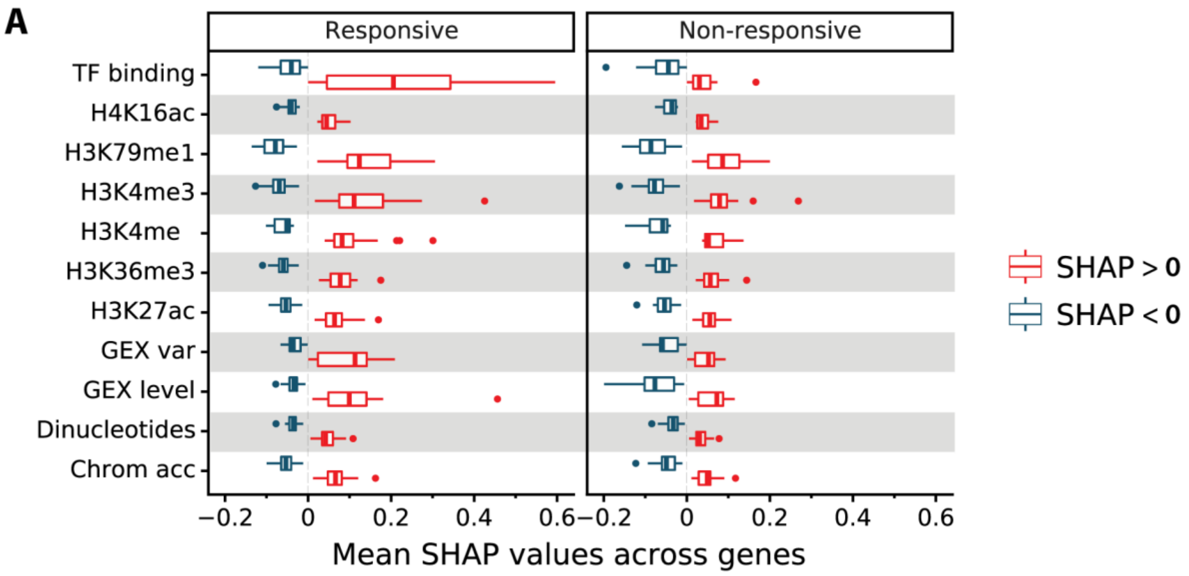


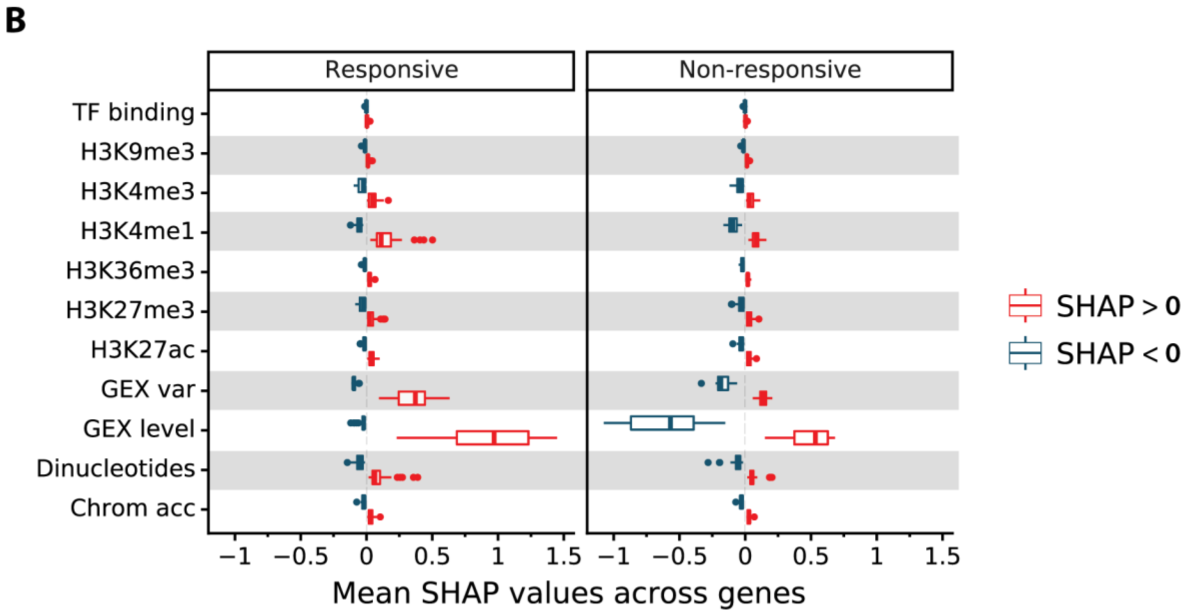


**Supplemental Figure S6.** (A) Mean SHAP values for all responsive and unresponsive targets of each yeast TF perturbation. (B) Mean SHAP values for all responsive and unresponsive targets of human TF perturbations.

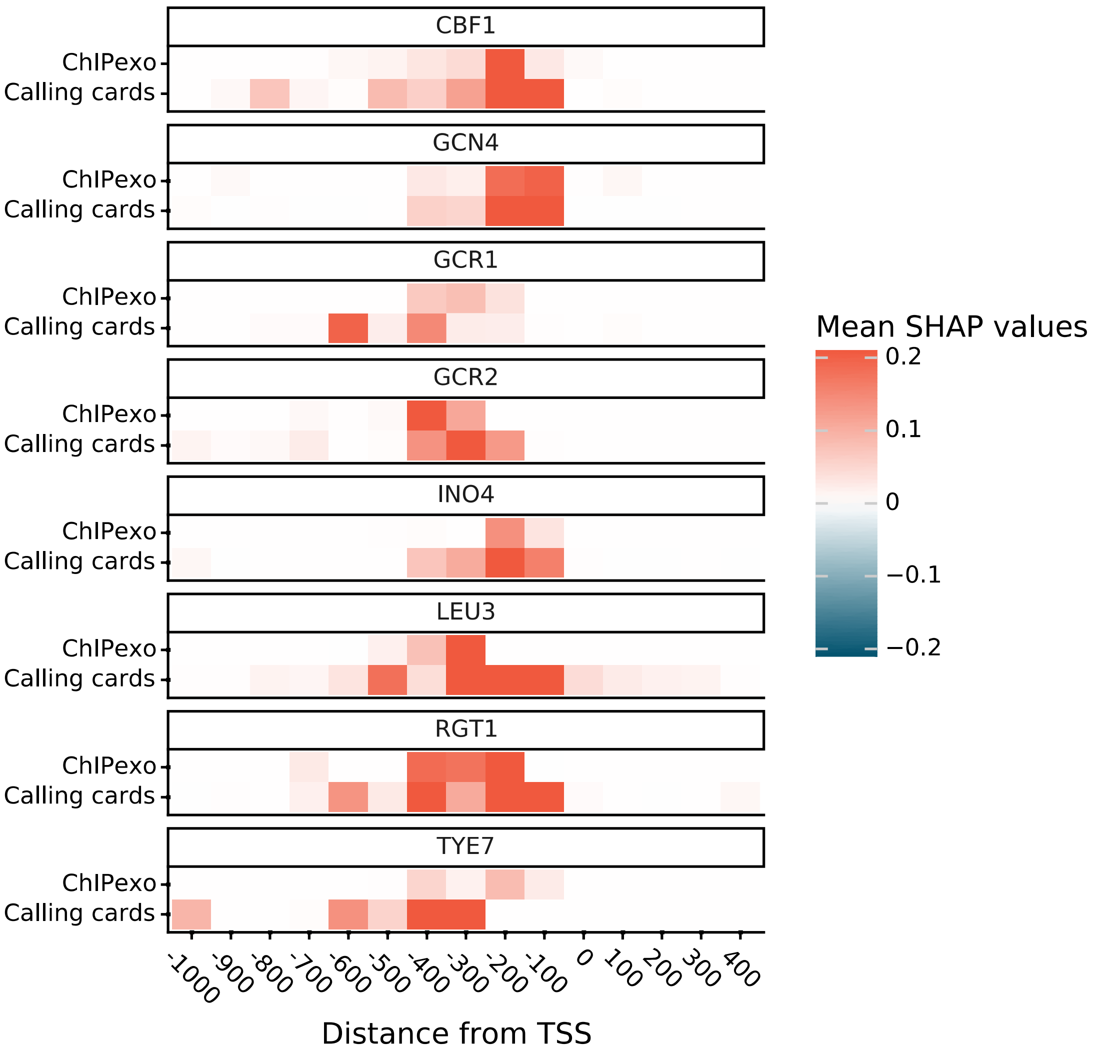


**Supplemental Figure S7.** Comparison of the influence of yeast TF binding data generated from two types of assays: transponson calling cards and ChIP-exo. Each pixel is the mean SHAP values of all target genes that were bound by the perturbed TFs.

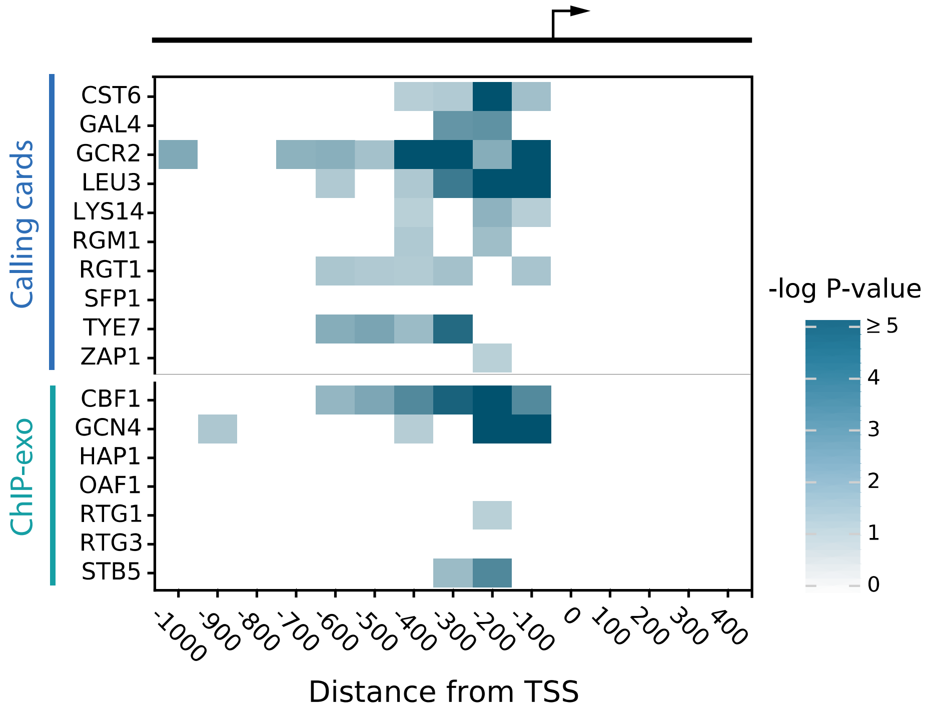


**Supplemental Figure S8.** Heatmap of the statistics that indicate the degree to which the bound but unresponsive genes have insufficient TF occupancy. The P-values for all bins in each row (TF) were estimated using the method in Figure 3C. Bins with P-values no less than 0.05 are in blank.

### SUPPLEMENTAL TABLES

#### Supplemental Table S1

| | Literatures | | | Data Availability | |
|---|---|---|---|---|---|
| | Karlić, 2010 | Zhou, 2014; González, 2015 | Kundaje, 2015; Singh, 2016 | Yeast (Weiner, 2015) | Human K562 (ENCODE, 2020) |
| H3K27ac | 1 | 1 | | 1 | 1 |
| H3K27me3 | | 1 | 1 | | 1 |
| H3K36me3 | 1 | | 1 | 1 | 1 |
| H3K4me1 | | 1 | 1 | 1 | 1 |
| H3K4me3 | 1 | 1 | 1 | 1 | 1 |
| H3K79me1 | 1 | | | 1 | |
| H3K9me3 | | | 1 | | 1 |
| H4K16ac | 1 | | | 1 | |

**Supplemental Table S2**

| TF | Binding dataset | AUPRC | For SHAP analysis |
|---|---|---|---|
| YLR403W (SFP1) | Calling cards | 0.58446718 | TRUE |
| YEL009C (GCN4) | ChIP-exo | 0.51278158 | TRUE |
| YLR451W (LEU3) | Calling cards | 0.36390632 | TRUE |
| YHR178W (STB5) | ChIP-exo | 0.3110723 | TRUE |
| YAL051W (OAF1) | ChIP-exo | 0.31106409 | TRUE |
| YIL036W (CST6) | Calling cards | 0.27532851 | TRUE |
| YNL199C (GCR2) | Calling cards | 0.27381191 | TRUE |
| YBL103C (RTG3) | ChIP-exo | 0.26980529 | TRUE |
| YLR256W (HAP1) | ChIP-exo | 0.24455603 | TRUE |
| YJR060W (CBF1) | ChIP-exo | 0.22821206 | TRUE |
| YDR034C (LYS14) | Calling cards | 0.21046638 | TRUE |
| YPL248C (GAL4) | Calling cards | 0.20618276 | TRUE |
| YJL056C (ZAP1) | Calling cards | 0.16135172 | TRUE |
| YKL038W (RGT1) | Calling cards | 0.16046672 | TRUE |
| YOR344C (TYE7) | Calling cards | 0.15421627 | TRUE |
| YMR182C (RGM1) | Calling cards | 0.12508439 | TRUE |
| YOL067C (RTG1) | ChIP-exo | 0.10535961 | TRUE |
| YEL009C (GCN4) | Calling cards | 0.45564519 | FALSE |
| YJR060W (CBF1) | Calling cards | 0.21908991 | FALSE |
| YNL199C (GCR2) | ChIP-exo | 0.21725465 | FALSE |
| YKL038W (RGT1) | ChIP-exo | 0.16007984 | FALSE |
| YLR451W (LEU3) | ChIP-exo | 0.1505335 | FALSE |
| YOR344C (TYE7) | ChIP-exo | 0.1503378 | FALSE |
| YOR363C (PIP2) | ChIP-exo | 0.09239194 | FALSE |
| YOL108C (INO4) | Calling cards | 0.07745282 | FALSE |
| YOL108C (INO4) | ChIP-exo | 0.06640978 | FALSE |

| | | | |
|---|---|---|---|
| YGL162W (SUT1) | ChIP-exo | 0.04185123 | FALSE |
| YMR280C (CAT8) | ChIP-exo | 0.03881355 | FALSE |
| YPL133C (RDS2) | ChIP-exo | 0.029646 | FALSE |
| YPL075W (GCR1) | ChIP-exo | 0.02614397 | FALSE |
| YPL075W (GCR1) | Calling cards | 0.02100747 | FALSE |
| YBR239C (ERT1) | ChIP-exo | 0.01386913 | FALSE |
| YJL089W (SIP4) | ChIP-exo | 0.01344086 | FALSE |

**Supplemental Table S3**

| TF | AUPRC | For SHAP analysis |
|---|---|---|
| ENSG00000105698 (USF2) | 0.64906387 | TRUE |
| ENSG00000074219 (TEAD2) | 0.63229787 | TRUE |
| ENSG00000131931 (THAP1) | 0.61089246 | TRUE |
| ENSG00000130522 (JUND) | 0.55512811 | TRUE |
| ENSG00000113658 (SMAD5) | 0.55354296 | TRUE |
| ENSG00000143390 (RFX5) | 0.54710206 | TRUE |
| ENSG00000177045 (SIX5) | 0.52840098 | TRUE |
| ENSG00000187098 (MITF) | 0.52106216 | TRUE |
| ENSG00000111206 (FOXM1) | 0.51931609 | TRUE |
| ENSG00000185591 (SP1) | 0.51506909 | TRUE |
| ENSG00000198176 (TFDP1) | 0.50122105 | TRUE |
| ENSG00000197905 (TEAD4) | 0.49858104 | TRUE |
| ENSG00000105722 (ERF) | 0.42194973 | TRUE |
| ENSG00000143379 (SETDB1) | 0.26877678 | TRUE |
| ENSG00000102145 (GATA1) | 0.25225519 | TRUE |
| ENSG00000141568 (FOXK2) | 0.24994403 | TRUE |
| ENSG00000144161 (ZC3H8) | 0.24814691 | TRUE |
| ENSG00000001167 (NFYA) | 0.22888833 | TRUE |
| ENSG00000134138 (MEIS2) | 0.21580418 | TRUE |
| ENSG00000186918 (ZNF395) | 0.2140645 | TRUE |
| ENSG00000162772 (ATF3) | 0.18328793 | TRUE |
| ENSG00000173039 (RELA) | 0.17614689 | TRUE |
| ENSG00000160633 (SAFB) | 0.17497082 | TRUE |
| ENSG00000123358 (NR4A1) | 0.14874691 | TRUE |
| ENSG00000115816 (CEBPZ) | 0.14337738 | TRUE |
| ENSG00000120837 (NFYB) | 0.14218726 | TRUE |

| | | |
|---|---|---|
| ENSG00000177463 (NR2C2) | 0.1354988 | TRUE |
| ENSG00000185551 (NR2F2) | 0.11380771 | TRUE |
| ENSG00000082641 (NFE2L1) | 0.11378921 | TRUE |
| ENSG00000172273 (HINFP) | 0.11147308 | TRUE |
| ENSG00000179348 (GATA2) | 0.09625769 | FALSE |
| ENSG00000060138 (YBX3) | 0.08859001 | FALSE |
| ENSG00000130254 (SAFB2) | 0.08360437 | FALSE |
| ENSG00000177485 (ZBTB33) | 0.07704929 | FALSE |
| ENSG00000126746 (ZNF384) | 0.07324795 | FALSE |
| ENSG00000112658 (SRF) | 0.06051329 | FALSE |
| ENSG00000158773 (USF1) | 0.05962755 | FALSE |
| ENSG00000126561 (STAT5A) | 0.05056387 | FALSE |
| ENSG00000106459 (NRF1) | 0.0484205 | FALSE |
| ENSG00000147421 (HMBOX1) | 0.00781738 | FALSE |
| ENSG00000156273 (BACH1) | 0.00700893 | FALSE |
| ENSG00000166478 (ZNF143) | 0.00116493 | FALSE |

### SUPPLEMENTAL METHODS

#### Data preparation

*TF-perturbation response data*

For yeast, we downloaded the microarray data for transcriptional responses 15 minutes after inductions of 194 TF using the ZEV induction system (Hackett et al. 2020). Column *log2_shrunken_timecourses* from the file “Raw & processed gene expression data” at https://idea.research.calicolabs.com/data was used as the levels of responses. Since these values were already shrunken towards zero during analysis, any gene with a non-zero value was defined as responsive.

For human, we used all RNA-seq expression profiles measured after gene knockout (KO) or knockdown (KD) in K562 cells from the ENCODE Project database (Dunham et al. 2012; Davis et al. 2018; Abascal et al. 2020). TFKO and TFKD mechanisms include TF-disabling mutations introduced by CRISPR, CRISPR inference (CRISPRi), small-interfering RNA (siRNA), and small-hairpin RNA (shRNA). We downloaded the expected counts of experimental and control profiles that were estimated using RSEM in the ENCODE RNA-seq pipeline and genome assembly GENCODE V24 (GRCh38). For each of the 355 experiments, we ran DESeq2 (V1.10.1) (Love et al. 2014) to identify differentially expressed genes by comparing the experimental replicates to the corresponding control replicates. Genes with Benjamini-Hochberg adjusted P-value < 0.05 and log2 fold-change > 0.5 were considered responsive.

*Pre-perturbation gene expression data*

The pre-perturbation expression level feature is the median gene expression level across all samples measured prior to the TF perturbation. For human genes (RNA-Seq data) we used the log TPM levels among all replicates of control samples. For yeast (microarray data) we used log fluorescence levels of the red (experimental) channel measured at time 0 (before each of the TF inductions). To construct a gene expression variation feature that is independent of the expression level, we used the method of (Sigalova et al. 2020). First, we computed the coefficient of variation (COV) of expression level in pre-perturbation samples for each gene. Next, we plotted all genes’ COV against their median expression level and fit a smooth curve using locally estimated scatterplot smoothing (LOESS) regression (Python scikit-misc V0.1.3). Each gene’s detrended expression variation feature was the residual of its COV from the LOESS fit (Supplemental Fig. S2: right panels).

*TF binding location data*

All coordinate-dependent features were mapped to yeast genome build sacCer3 and human build GRCh38. Yeast binding location data were generated using transposon calling cards (Wang et al. 2011; Shively et al. 2019; Kang et al. 2020) or ChIP-exo (Bergenholm et al. 2018; Holland et al. 2019). For the calling cards data (16 TFs) we lifted over the transposon insertion coordinates, which were originally mapped based on sacCer2, to sacCer3 using the LiftOver tool in UCSC genome browser. ChIP-exo peaks for 20 yeast TFs were obtained directly from the authors of Bergenholm et al. (2018) and Holland et al. (2019). Kang et al. (2020) reported that among the four environmental conditions, the bound targets of these TFs in glucose limited chemostat condition have the best agreement with the perturbation-responsive targets at 15 minutes after TF induction, so we focused on this condition. Peak locations were reported for the genome of yeast strain CEN.PK, so we lifted them over to coordinates in sacCer3 (strain S288C) as follows. First, since the loci of TFBSs were reported as relative distances to CEN.PK TSSs, these loci were converted to the relative distances to the ORFs using the CEN.PK TSS annotation (https://github.com/SysBioChalmers/ChIPexo_Pipeline/blob/master/Data/TSSData.tsv). Due to high similarity of the two yeast strains, we assumed that the relative distance of each TFBS to CEN.PK ORF are the same for the matching S288C ORF. Next, the relative distances were converted to absolute genomic coordinates in sacCer3 using S288C gene annotation from the *Saccharomyces* Genome Database (SGD).

We downloaded ChIP-seq peaks for human 54 TFs in K562 cells from the ENCODE Project (Dunham et al. 2012; Davis et al. 2018; Abascal et al. 2020). K562 had far more TFs that were both ChIPped and perturbed than any other cell type. We only consider gene to be a TF if has a well-defined DNA-binding factors according to ref. (Lambert et al. 2018). We used only the "conservative" peaks, which had Irreproducible Discovery Rate <= 2%. The log10 q-value reported for each peak was used as the binding signal feature.

*Histone modifications and chromatin accessibility data*

For yeast, we used histone modification data harvested prior to the addition of a diamide stress from ref. (Weiner et al. 2015), which was produced using MNase-ChIP-Seq (GEO accession GSE61888, https://www.ncbi.nlm.nih.gov/geo/). We used chromatin accessibility data at harvested prior to the introduction of osmotic stress from ref. (Schep et al. 2015) (GSE66386). For human K562 cells, we downloaded the coverage data (fold change over control) for histone modifications and chromatin accessibility from ENCODE (Dunham et al. 2012; Davis et al. 2018; Abascal et al. 2020).

*Mapping genome-wide features to cis-regulatory regions*

We defined yeast promoter regions as 1,000 bp upstream to 500 bp downstream from the transcription start site (TSS). TSS coordinates were obtained from ref. (de Boer et al. 2020). Inputs for each genome-wide feature were mapped relative to the TSS and summed over each of 15 100-bp-bins to create 15 features.

For the human, TSS coordinates were downloaded from Ensembl Release 92 (Cunningham et al. 2019). For each gene, we defined the *5' promoter* to be 4 Kb centered on the 5'-end TSS, *alternative promoters* to be 4 Kb region centered at any TSS that is more than 2 Kb from the 5'-

end TSS, and *enhancers* to be enhancers that are linked to the gene in the GeneHancer V4.8 database (Fishilevich et al. 2017). We used only “double elite” enhancers, meaning that the enhancers existence and linkage to the target gene are both supported by at least two distinct types of evidence. Enhancers that are more than 500 Kb from the 5’-end TSS were removed.

Each gene must have an equal number of features to create a rectangular feature matrix, even though genes differ greatly in how many alternative TSSs and enhancers they have. For TSSs, we primarily focused on 5’-end, for two reasons. First, approximately half of all alternative TSSs fall within the 2 Kb region downstream from the 5’-end TSS (Supplemental Fig. S1A). The promoter regions of these nearby alternative TSSs largely overlap with that of the 5’ TSS, so we did not think it necessary to create separate features for them. Second, the 5’ TSS and others within 2 kb of it account for approximately three times as much transcription as the TSSs usage outside the region, according to Fantom5 CAGE data (Forrest et al. 2014; Lizio et al. 2019) (Supplemental Fig. S1B). Alternative promoters outside of this region (median distance 26 kb) were treated as enhancers, since enhancers and promoters share most of their properties and functions (Andersson and Sandelin 2020). Signals located in enhancers were aggregated into features in two ways. The first method (*Prom + bin enhan*; Fig. S3, blue) sums signals within each of 45 bins upstream of the 5’ end promoter region and another 45 bins downstream. The width of the bin closest to the TSS was 500 bp and each subsequent bin was larger by 500 bp. For example, the widths of the three bins closest to the TSS are 500, 1000, and 1500 bp. Together, the 45 bins on each side of the promoter covered 498 Kb (-500 Kb to -2 Kb or +2 Kb to +500 Kb). The second method (*Prom + agg enhan*, Fig. S3, green) sums enhancer signals 500 Kb upstream or downstream of the promoter into two features. Note that only signals that fell within defined enhancers were used.

#### Predicting TF-perturbation responses using cross-validation

For each TF perturbation, we trained and tested a model for predicting whether a gene will respond by using 10-fold cross-validation. The genes in each fold were selected at random subject to the constraint that all folds have the same proportion of responsive genes. We tried two ensemble classification algorithms—random forests implemented in scikit-learn library (V0.22.1) (Pedregosa et al. 2011), and gradient boosted trees implemented in XGBoost library (V 0.90) (Chen and Guestrin 2016). For both methods, 500 trees were used. For XGBoost, learning rate of 0.01 was used for the “gbtree” booster. All other parameters were default. In each cross-validation fold, training data were transformed to Z-scores and test data were consequently transformed by using the scaling factors learned from training data. Precision-recall curves were calculated by comparing the predicted probability of response to the true categorization of genes as responsive or responsive.

#### Using SHAP to quantify the influences of features on predictions

We employed the SHAP (SHapley Additive exPlanations) framework (V0.35.0) (Lundberg and Lee 2017) to quantify the extent to which each feature contributes to the predicted probability of responsiveness for each gene (TreeExplainer function). SHAP uses a linear model to approximate the predictions for artificially constructed instances in the neighborhood of each actual instance. SHAP values explain why one particular prediction – one TF-gene pair – differs from the average prediction for all genes in response to perturbation of that TF. Positive values indicate how strongly a particular feature value pushes the model toward assigning the gene a higher probability of responding, while negative values represent how strongly the value pushes the model toward assigning the gene a lower probability of responding.

#### Aggregating SHAP values across genes sets

To characterize the effect of a particular feature for a set of genes, we separately summed its positive and negative SHAP values over genomic bins for each gene. We then averaged the positive sums over all genes and, separately, the negative sums. Formally, we calculated mean positive SHAP value $S_k^+$ and mean negative SHAP value $S_k^-$ as:

$$S_k^+ = \sum_{i \in G'} \sum_{j \in B} \phi_{ijk} \, \mathbb{1}_{(0,+\infty)}(\phi_{ijk}) / |G'|$$

$$S_k^- = \sum_{i \in G'} \sum_{j \in B} \phi_{ijk} \, \mathbb{1}_{(-\infty,0)}(\phi_{ijk}) / |G'|$$

where $\phi_{ijk}$ is the SHAP value for gene $i$ in bin $j$ for feature $k$, $G'$ is the set of gene indices, ($G' \subseteq$ G, where $G$ is for all genes), and $B$ is the set of bin indices. For coordinate-independent features (e.g., pre-perturbation gene expression), $B$ has size of one. We defined the *Net influence* of a feature on predictions for a set of genes as a single sum including both positive and negative SHAP values of the feature over the set of genes. *Net influence* measures the feature's overall direction of influence. *Global feature importance* is the sum of absolute values of the SHAP values of a feature for genes in the set. *Global feature importance* measures how important the feature is in determining the model's prediction, regardless of direction.

#### Modeling and interpreting generic responses in any genetic perturbation

We used an XGBoost regressor to predict the fraction of genetic perturbations to which each gene responds, in absence of perturbation-specific information. Three data sets consisted of all TF induction samples in yeast (n=194), all TFKO/TFKD samples in K562 cells (n=56), and all genetic perturbations in K562 (n=355). The corresponding feature matrix for each label vector includes all features except for TF binding data. Performance was evaluated by cross validation as described above.
